# Discovery of EMRE in fungi resolves the true evolutionary history of the mitochondrial calcium uniporter

**DOI:** 10.1101/2020.03.24.006015

**Authors:** Alexandros A. Pittis, Valerie Goh, Alberto Cebrian-Serrano, Jennifer Wettmarshausen, Fabiana Perocchi, Toni Gabaldón

## Abstract

Mitochondrial calcium (mt-Ca^2+^) uptake is central for the regulation of numerous cellular processes in eukaryotes^1^. This occurs through a highly selective Ca^2+^ uniporter located at the inner mitochondrial membrane and driven by the membrane potential^2–4^. While the physiological role of the uniporter was extensively studied for decades, its genetic identity was only recently determined, with MCU^5,6^, MICU1^7^ and EMRE^8^ constituting pore-forming and regulatory subunits. Preliminary evolutionary analyses suggested an ancient eukaryotic origin of mt-Ca^2+^ uptake, but also pinpointed inconsistent phylogenetic distributions of MCU, MICU1, and EMRE within fungi, where homologs of MCU were present in the absence of the supposedly essential regulators, MICU1 and EMRE^9,10^. Here, we perform the most comprehensive phylogenomic analysis of the mt-Ca^2+^ uptake system and trace its evolution across 1,156 fully-sequenced eukaryotes. In contrast to earlier assumptions^9–11^ we find compelling evidence that previously identified animal and fungal MCUs, the targets of several structural and functional efforts^11–16^, represent two distinct paralogous subfamilies originating from an ancestral duplication. We further uncover a complete “animal-like” uniporter complex within chytrid fungi, including bona-fide orthologs of MCU, MICU1, and EMRE. This first identification of EMRE outside Holozoa (animals and their unicellular relatives) and its strong coevolution with “animal-like” MICU1 and MCU indicates that these three components formed the core of the ancestral opisthokont uniporter. We confirm this finding experimentally, by showing that chytrid EMRE orthologs in combination with either human or “animal-like” MCUs, but not with “fungal-specific” MCUs, can reconstitute mt-Ca^2+^ uptake *in vivo* in the yeast *Saccharomyces cerevisiae*. Hence, we here solve a purported evolutionary paradox: the presence of MCU homologs in fungal species devoid of other uniporter components and with no detectable mt-Ca^2+^ uptake. Altogether, our study clarifies the evolution of the mt-Ca^2+^ uniporter and identifies new important targets for comparative structural and functional studies.

## Main text

Comparative genomics analyses, based on a few eukaryotic species, in combination with RNAi assays were instrumental in the identification of MCU and MICU1 as the founding members of the mt-Ca^2+^ uniporter^5–7^. This discovery paved the way to the identification of other paralogous components of this channel in mammalian cells, including negative and positive regulatory and tissue-specific subunits (MCUb^17^, MICU2^18,19^, MICU3^20^, and MICU1.1^21^). Instead, EMRE was found as a specific interactor of MCU in human cells, required for both conductivity and binding of the channel to MICU1^8^. Although MCU and MICU1 showed correlated evolutionary histories across 138 sequenced eukaryotic organisms, EMRE apparently lacked any homolog outside the metazoan lineage and it was therefore suggested to be an animal-specific innovation^9–11^. While those observations pointed to an ancient eukaryotic origin of mt-Ca^2+^ uptake, they also implied a very different composition and regulation of the uniporter in different clades. A notable case is the identification of MCU as the only uniporter component in Basidiomycota and filamentous Ascomycota (e.g., *Neurospora crassa* and *Aspergillus fumigatus*), suggesting that either fungal MCUs are sufficient for mt-Ca^2+^ uptake or they are regulated independently of MICU1 and EMRE^9,10^. Based on the assumption of an orthologous relationship between human and fungal MCUs^9,10,22^, several independent structural studies of Ascomycota MCUs have been performed to understand the basic principles of uniporter channel assembly and function^12–15^. However, those organisms had been shown to lack uniporter activity^23,24^ and their MCU homologs were unable to mediate mt-Ca^2+^ uptake when heterologously expressed in HeLa or yeast cells^14,25^. Not surprisingly, significant structural and sequence differences were found between fungal MCUs and their animal counterparts^12–15^, raising the question of whether fungal MCUs function as classical Ca^2+^ uniporters at all.

To resolve this paradox, we assessed the evolution of each uniporter component across 1,156 fully-sequenced eukaryotic genomes (see **Supplementary Table 1**), using a combination of profile-based sequence searches, protein domain composition assessment, and phylogenetics. As shown in **Fig. 1** (**Extended Data Fig. 1**), the overall taxonomic distributions of MCU and MICU1 homologs were largely congruent with that of previous genomics surveys^9,10^. We confirmed the presence of MCU in at least some species of the major eukaryotic groups (Unikonts, the SAR clade, Plants and Euglenozoa), and its absence in all sequenced Apicomplexans, all yeasts in Saccharomycotina and most in Schizosaccharomyces clades, Microsporidia, *Trichomonas* and *Giardia*. Hence, mt-Ca^2+^ uptake appeared to have been lost many times independently during the evolution of eukaryotes. A significant number of these losses correlated with extreme streamlining of mitochondrial metabolism, as most MCU/MICU-lacking lineages encompassed relict forms of anaerobic mitochondria, such as mitosomes (Microsporidians, *Entamoeba, Giardia, Cryptosporidium*) or hydrogenosomes (*Trichomonas*)^26^. Our homology-based results confirmed the above-mentioned anomaly that most fungal genomes, for which our dataset is particularly rich - 776 species as compared to 50 in previous studies^9,10^ – encode for MCU but not MICU or EMRE. Unexpectedly, however, our analysis uncovered for the first time the presence of EMRE outside Holozoa, identifying reliable orthologs in three chytrid fungi – an early-diverging zoosporic fungal lineage: *Allomyces macrogynus, Catenaria anguillulae*, and *Spizellomyces punctatus*. Additional searches in public databases confirmed that EMRE was not present in other sequenced fungi.

**Figure 1:**
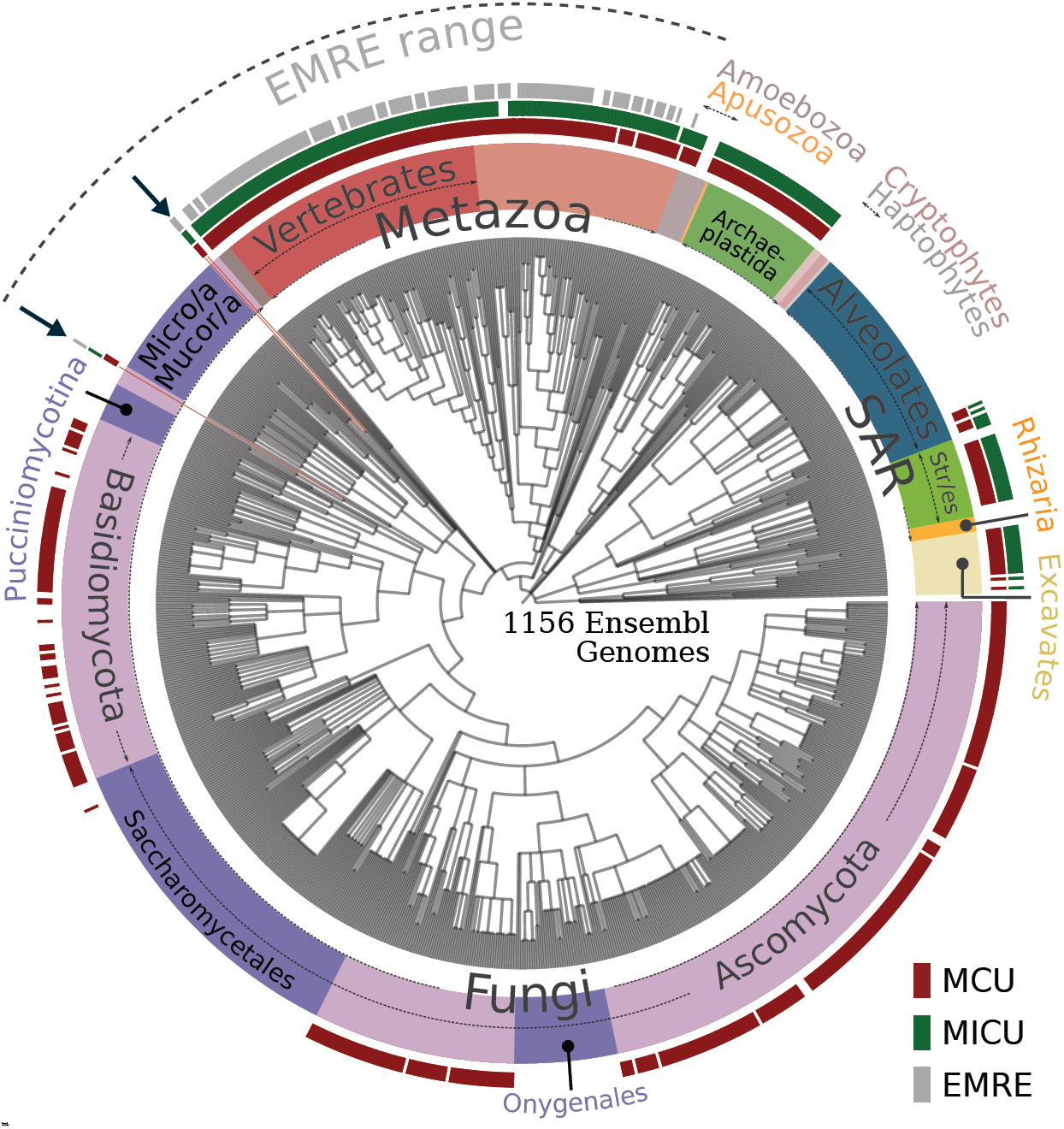
Phylogenetic distribution of mt-Ca^2+^ uniporter protein families. The phylogenetic distribution of MCU (red), MICU1 (green), and EMRE (gray) homologs across 1,156 eukaryotic genomes is shown on the NCBI taxonomy tree. Viridiplantae and Rhodophyta (red algae) have been grouped together as Archaeplastida, and Alveolates, Stramenopiles (Str/es) and Rhizaria as the SAR clade. In all cases where data from various strains of a species are present with the same pattern, these have all been collapsed to the species level, resulting in 969 terminal nodes shown. The mt-Ca^2+^ uniporter complex has been completely lost in Apicomplexa within Alveolates, Rhizaria (5 genomes), red algae (3 genomes), Cryptophytes (3 genomes), Haptophytes (1 genome), and the Entamoeba clade within Amoebozoa. Within fungi (in purple), all major clades that have completely lost MCU homologs are indicated with a darker purple color, namely Onygenales, Saccharomycetales, Pucciniomycotina, Mucoromycotina (Mucor/a), and Microsporidia (Micro/a). The only three early diverging fungal species (*A.macrogynus, C.anguillae, S.punctatus*) that encode also MICU and EMRE are highlighted with a red arrow. The NCBI taxonomy and the presence/absence profile were visualized using the ETE toolkit^33^. For a version of the profile, which includes the species names, see Extended Data Figure 1.

To clarify the underlying evolutionary history of the uniporter, we reconstructed and inspected the molecular phylogenies of MCU (**Fig. 2a, Extended Data Fig. 2a**) and MICU1 (**Fig. 2b, Extended Data Fig. 2b**) homologs across eukaryotes. We found that the evolution of both MCU and MICU gene families was driven by numerous gene duplications and losses, some of them having occurred in parallel in different lineages, implying an ancient and tight functional relationship. Furthermore, we showed that evolutionary independent duplications at the base of several main eukaryotic lineages – Vertebrates, Streptophytes, Oomycetes, Kinetoplastids, and Ciliates – resulted in the existence of multiple MCU paralogous copies in each of these clades. As a result, orthology relationships within the MCU gene family are complex and of the type many-to-many^27^. This means, for instance, that human MCU and MCUb are equally distant evolutionarily (co-orthologous) to each non-vertebrate animal MCUs, and also to all MCUs from protists and plants, explaining why *Trypanosoma*’s MCUb does not functionally complement human MCUb^28,29^. It also implies that shared physical interactions between paralogous MCU proteins in hetero-oligomers in different organisms, as shown for human and *Trypanosoma brucei* MCUb proteins^17,28,29^, are the result of parallel evolution. Importantly, one duplication event in the MCU gene family occurred in the common ancestor of opisthokonts and was followed by differential losses that distinguished Holomycota (fungi and their relatives, including *Fonticula alba*) from the other opisthokonts, i.e. the Holozoa (animals and their unicellular close relatives). Most fungal species kept only one of the two MCU paralogs that is referred here as the “fungal-specific” MCU (fsMCU). Holozoa, instead, retained the other MCU paralog, the bona-fide “animal” MCU. Only three chytrid fungi in our dataset, *A. macrogynus, C. anguillulae*, and *S. punctatus* retained both the fsMCU and the “animal” MCU, and these are also the only fungi encoding MICU1 and EMRE homologs. This striking, previously undetected, co-evolution pattern between MCU, MICU and EMRE in fungi suggests a strong interdependence, and even stronger considering that Blastocladiales (*Allomyces* and *Catenaria*) and *Spizellomyces* do not form a monophyletic clade^30^. Based on these findings, we hypothesized that these “animal-like” MCUs present in chytrids should require EMRE to drive mt-Ca^2+^ uptake, similarly to their human ortholog.

**Figure 2:**
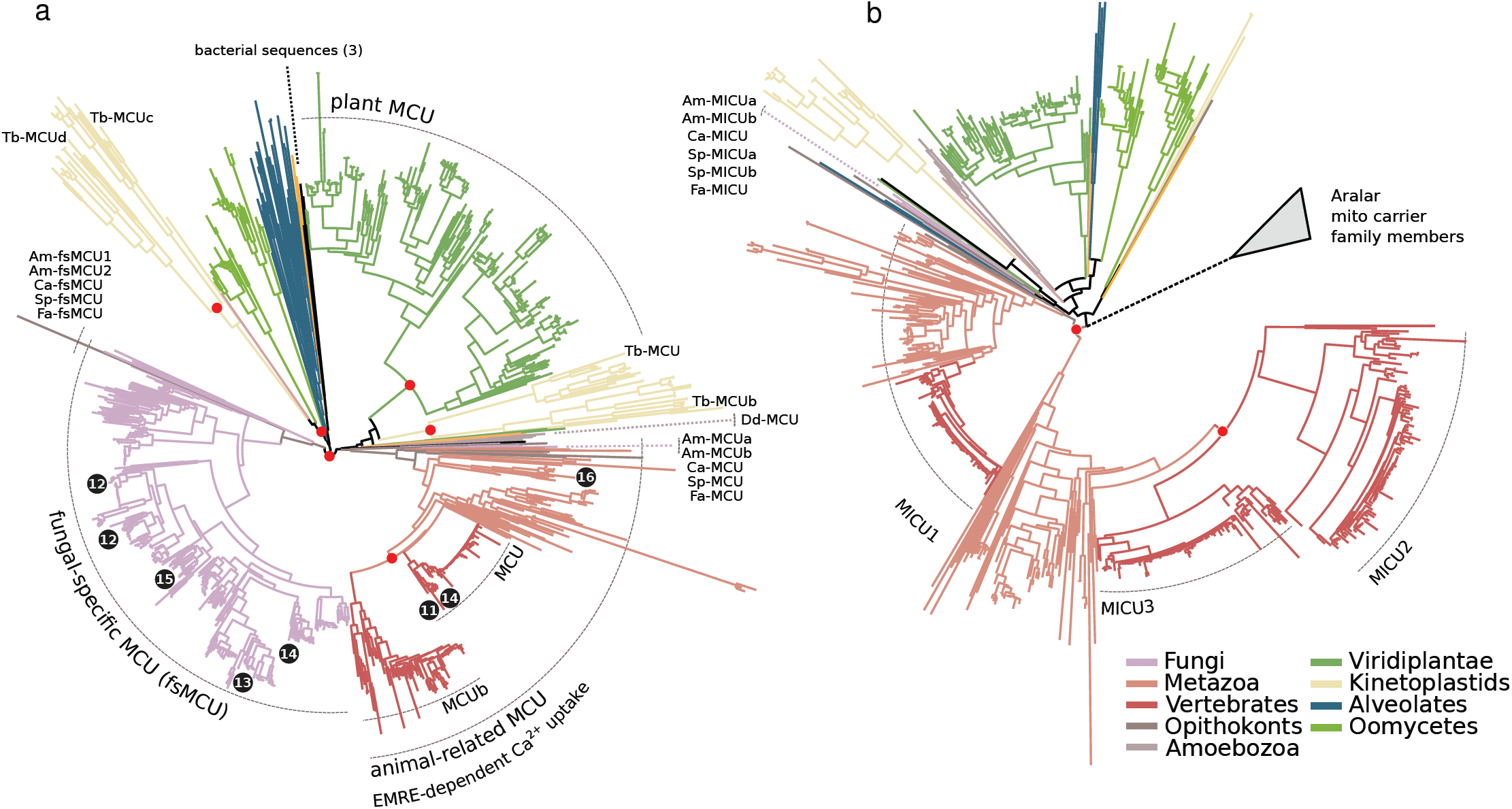
The fungal-specific MCU sub-family is a distinct phylogenetic clade. Maximum Likelihood phylogenetic trees of MCU (**a**) and MICU (**b**) families. The two families have been expanded through duplication rounds in various independent lineages. Major duplication events are indicated with a red sphere on the relevant tree node. In both (**a**) and (**b**) names and relative positions in the trees of members from Holomycota and few other representative species are shown. The main sub-families (MCU/MCUb and MICU1/2/3) were named after the human/mouse sequences within the phylogenetic clades. The phylogenetic positions of MCU orthologs with published structural data are shown in black circles, while the numbers refer to the publication reference. fsMCU, “fungal-specific” MCU. In (**b**) all Aralar-related MICU homologs were excluded after a first pre-processing, but are shown here as reference (See also Methods).

The above mentioned finding of bona-fide EMRE orthologs in all these three cythrids (**Fig. 1**) reinforced this idea and placed back the origin of an animal-like mt-Ca^2+^ uptake in the opisthokont ancestor, preceding the diversification of animals and fungi. Consistently, the heterologous expression of MCU from *Dictyostelium discoideum*, representing an amoebozoan lineage that diverged earlier than the origin of opisthokonts, is alone sufficient to reconstitute mt-Ca^2+^ uptake in yeast mitochondria, while human (Hs-) MCU only does so in the presence of EMRE^31^. Similarly, we hypothesized that co-expression of “animal” MCUs and EMRE proteins from chytrids would be necessary and sufficient to reconstitute uniporter activity. The phylogenetic distribution profile (presence/absence) across the MCU complex components reveals a strong co-evolution pattern, when only the true orthologous sequences are considered (**Fig. 3a**). Strikingly, we detected mt-Ca^2+^ uptake in yeast strains expressing “animal” MCUs from either *A. macrogynus* (Am-MCUa) or *S. punctatus* (Sp-MCU) with their respective EMREs (Am-EMRE, Sp-EMRE) (**Fig. 3b,c, Extended Data Fig. 3**). In contrast, we did not detect any mt-Ca^2+^ uptake in yeast strains expressing fsMCU proteins from *A. macrogynus* (Am-fsMCU1) and *S.punctatus* (Sp-fsMCU), despite proper expression and localization (**Extended Data Fig. 3** and **Extended Data Fig. 4**). Similar results would be expected in other Holozoa despite the inability to detect EMRE by similarity searches. Indeed, the co-expression of MCU from the sea anemone *Nematostella vectensis* (Nv-MCU) with Hs-EMRE in yeast was able to reconstitute mt-Ca^2+^ uptake to a similar extent of a strain expressing Hs-MCU and Hs-EMRE (**Extended Data Fig. 5**). These results, together with the absence of MICU proteins in most fungal lineages, indicate that mt-Ca^2+^ uptake in fungal mitochondria, if it exists, is not mediated by fsMCUs, or that a different –yet unknown-regulator is necessary. Instead, animal-like MCUs from chytrid fungi and Holozoa function similarly to the mammalian uniporter, in an EMRE-dependent fashion. Altogether, our evolutionary analyses and experimental results confirm that MCU-EMRE interaction is conserved, and was already present in the last common ancestor of fungi and animals.

**Figure 3:**
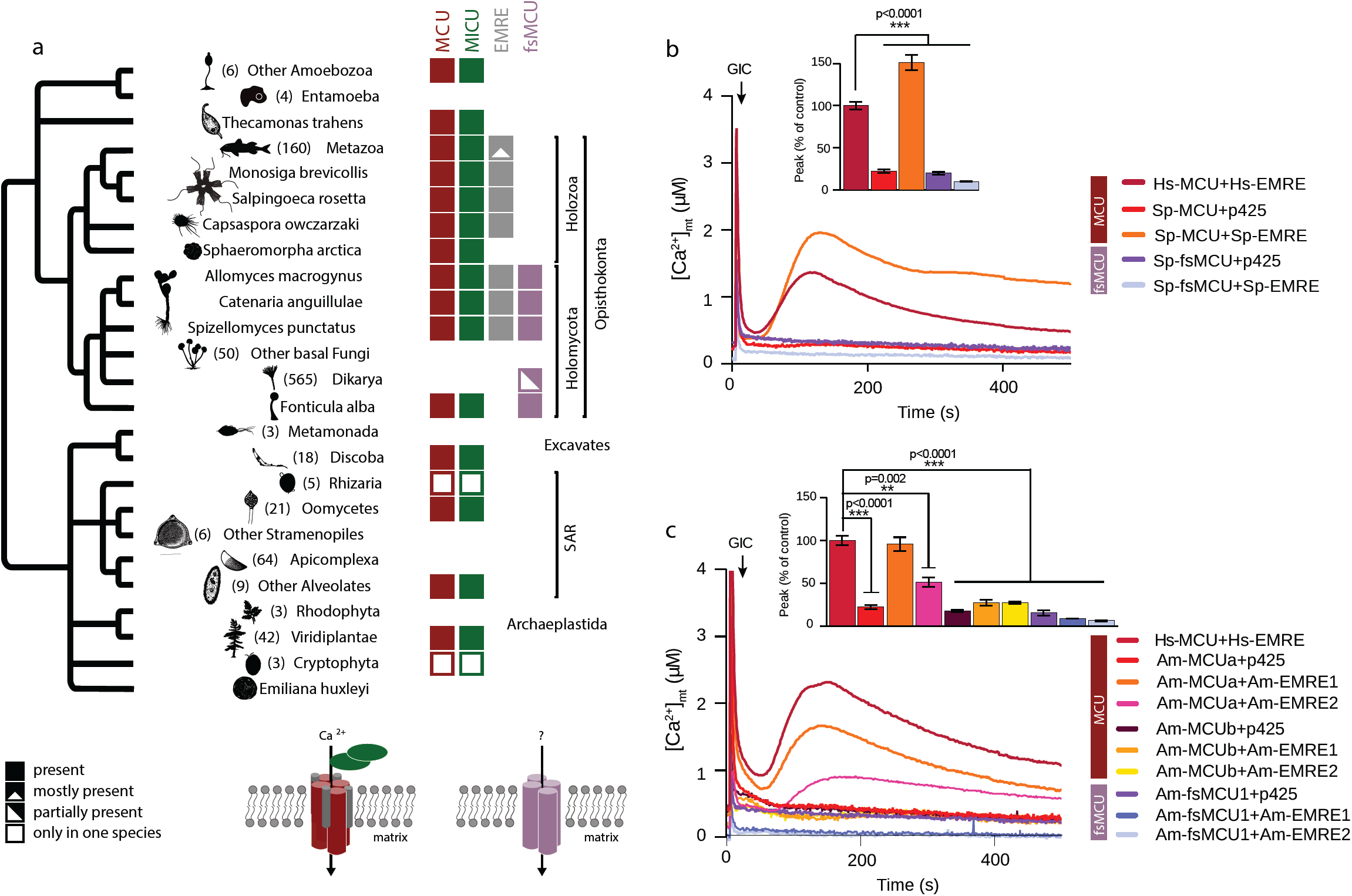
Functional reconstitution of mt-Ca^2+^ uptake by fungal MCU and EMRE orthologs. **a**, Phylogenetic distribution profile (presence/absence) across MCU complex components. The distribution pattern of MICU and EMRE largely overlaps with that of the animal like MCU, but not the fsMCU. **b,c**, Representative traces and quantification of mt-Ca^2+^ transients in yeast cells expressing animal-like or fungal-specific MCU orthologs from S. *punctatus* (**b**) and *A. macrogynus* (**c**) with either their respective EMRE proteins or empty vector (p425) upon glucose-induced calcium (GIC) stimulation in presence of 1 mM CaCl_2_. All data represent mean ± SEM (n=3); ***p < 0.0001, **p < 0.001, one-way ANOVA with Dunnett’s Multiple Comparisons Test.

Consistent with our hypothesis that fsMCUs do not represent true functional orthologs of Hs-MCU, when comparing MCU sequences across eukaryotes we found that fsMCUs lack key residues conserved in the animal-like MCUs, despite retaining a DXXE motif (**Fig. 4a** and **Extended Data Fig. 6a**). Those residues have been previously shown to be important for MCU function and its interaction with EMRE^32^, Notably, animal and fungal EMREs appear highly divergent (**Fig. 4b** and **Extended Fig. 6b**), although the MCU interacting domain GXXXA/S/G and the polyaspartate tail necessary for the binding to MICU1^32^ are fully conserved. Interestingly, the fungal EMRE sequences contained an extra C-terminal domain that is not found in Holozoa, suggesting some degree of specialization. Thus, we hypothesized that Am-MCUa and Sp-MCU would have evolved to interact with EMRE proteins from the same or related species. Indeed, those animal-like MCUs were neither able to increase mt-Ca^2+^ uptake when expressed in a wild-type HeLa background nor to rescue mt-Ca^2+^ uptake in MCU knock-down (shMCU) HeLa cells (**Extended Data Fig. 7**), despite showing proper expression and insertion into the inner mitochondrial membrane (**Extended Data Fig. 8**). Furthermore, the expression of Am-MCUa and Sp-MCU in yeast mitochondria was unable to reconstitute mt-Ca^2+^ uptake in the presence of Hs-EMRE (**Fig. 4c** and **Extended Data Fig. 9a**). Instead, Hs-MCU was functional when co-expressed with either Am-EMRE or Sp-EMRE (**Fig. 4d** and **Extended Data Fig. 9b**). Altogether, these results suggest that the C-terminal domain of fungal EMREs is dispensable for a functional interaction with Hs-MCU but necessary to activate animal-like fungal MCUs. Accordingly, we observed that the co-expression of Am-MCUa and Sp-MCU with Am-EMRE and Sp-EMRE lacking the extra C-terminal domain (EMRE-t) was unable to efficiently reconstitute mt-Ca^2+^ uptake (**Fig. 4e** and **Extended Data Fig. 10a**). However, while the presence of the extra C-terminal domain in Hs-EMRE did not affect the function of Hs-MCU, it was not sufficient to reconstitute mt-Ca^2+^ uptake when co-expressed with Am-MCUa and Sp-MCU (**Fig. 4f** and **Extended Data Fig. 10b**). On the one hand, these findings hint a possible activating role of the extra C-terminal domain of the fungal EMRE for the efficient regulation of animal-like MCUs function. On the other hand, they suggest that functional domains that are conserved between fungal and human EMREs are sufficient to regulate Hs-MCU activity.

**Figure 4:**
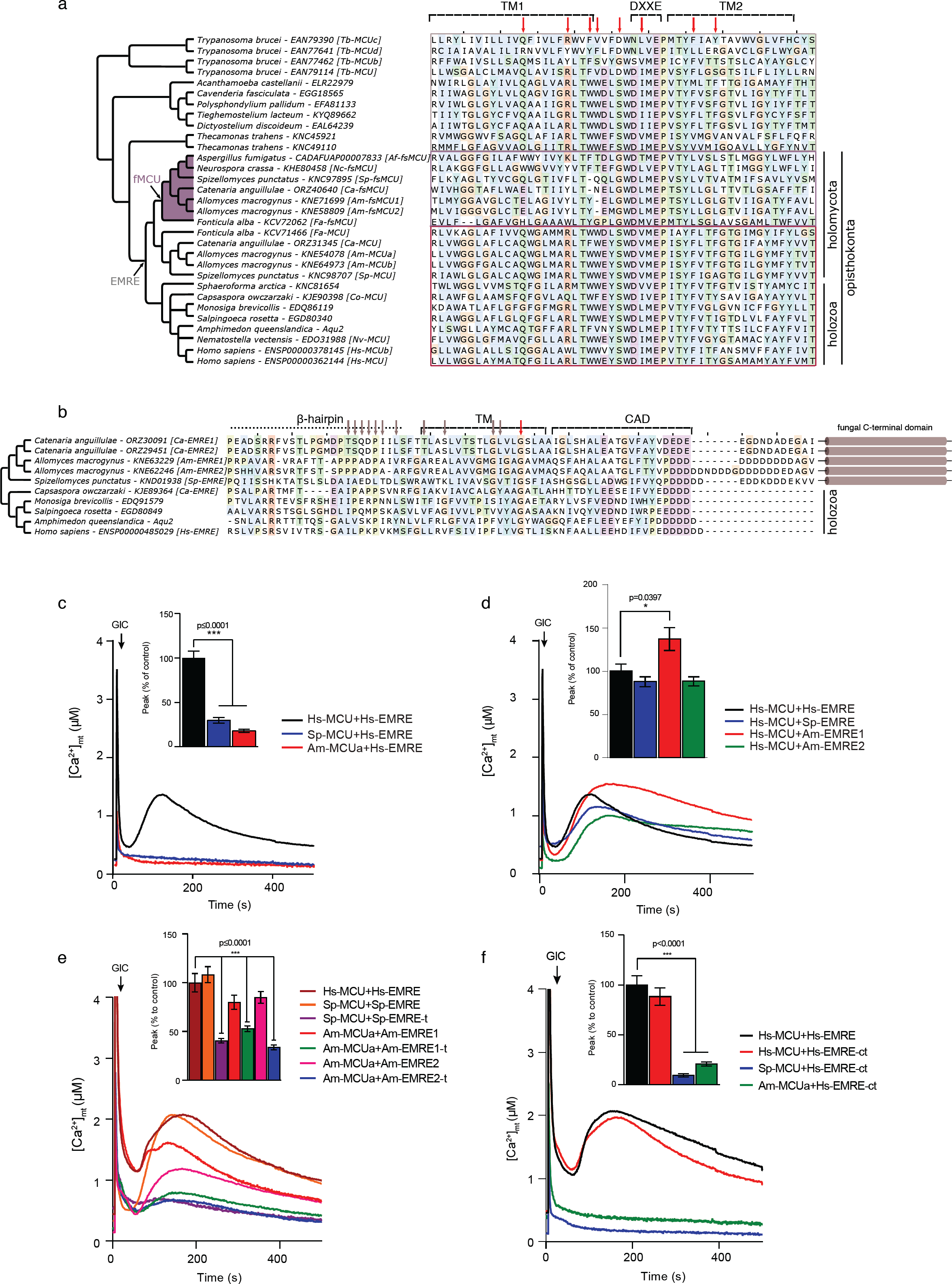
Evolution of MCU-EMRE interaction. **a,b**, Phylogenetic trees of members of MCU (**a**) and EMRE (**b**) protein families, and sequence diversity of major domains. The sequence alignment of TM1 and TM2 of MCU sequences from 20 species is shown in (**a**). The program Multi-Harmony^34^ was used to detect residues that are overall conserved but differ in the fsMCU members (highest scoring positions are indicated with red arrows). The fsMCU clade is shown in purple. The evolutionary point where the MCU proteins become EMRE-dependent is shown in gray. The degree of conservation across the animal related MCU members is very high in these loci, while few positions are Holomycota or Holozoa specific. Similarly in (**b**) EMRE’s sequence diversity across opisthokonts is shown for the β-hairpin, the TM, and CAD domain. Residues found important for the interaction between MCU and EMRE in^11^ and fully conserved positions are indicated with gray and red arrows, respectively. **c,d,e,f** Representative traces and quantification of mt-Ca^2+^ transients in yeast cells expressing either human and animal-like *S. punctatus* and *A. macrogynus* MCU with human EMRE (**c**), species-specific EMRE with human MCU (**d**), animal-like S. *punctatus* and *A. macrogynus* MCU with their respective wild-type or truncated (-t) EMREs (**e**) or with human EMRE fused to the fungal extra C-terminal domain (**f**) upon glucose-induced calcium (GIC) stimulation in presence of 1 mM CaCl_2_. All data represent mean ± SEM; n=4; ***p < 0.0001, *p < 0.01, one-way ANOVA with Dunnett’s Multiple Comparisons Test.

Altogether, our identification of a strong co-evolution pattern between MCU, MICU and EMRE provides an explanation to an elusive evolutionary paradox: the presence of uniporter homologs in species with no detectable mt-Ca^2+^ uptake^9,10^. We demonstrate that an animal-related MCU complex has been lost early within the evolution of most fungal clades, which retained only fungal-specific paralogous MCUs protein (fsMCUs). These fsMCUS, which are paralogous to Hs-MCU, have been so far wrongly considered functionally equivalent to Hs-MCU and deeply studied for this reason. This is the case, for instance, of the Ascomycota MCUs proteins (e.g., *Neurospora crassa*) that have been recently used as models for understanding structure and regulation of the human uniporter channel^11–16^. Consistently, we and others find that fungal MCUs are unable to reconstitute or rescue mt-Ca^2+^ uptake in yeast or HeLa cells lacking MCU, respectively^14,23,25^. Similarly, the same fungal MCU sequences used for investigating structure and function of the mammalian uniporter^12–14^ failed to mediate mt-Ca^2+^ uptake when expressed in yeast mitochondria or in MCU knockout HeLa mtAEQ cells (**Extended Data Fig. 11**). Instead, we show that only fungal species having both MCU and EMRE sequences (animal-like MCUs), such as *A. macrogynus* and *S.punctatus*, are able to mediate mt-Ca^2+^ uptake. Those observations together with the significant structural and sequence differences found between fungal and animal MCUs^11^ question whether fungal MCUs function as classical Ca^2+^ uniporters.

We identify, for the first time, non-metazoan EMRE sequences and demonstrate the ancestrally essential role of EMRE in mt-Ca^2+^ homeostasis. Our results imply that EMRE, previously thought to be an animal-specific innovation^8^, formed part of an animal-like machinery in the common ancestor of opisthokonts, and was lost secondarily in the evolution of fungi together with the other components of the animal-like mt-Ca^2+^ uptake machinery (**Fig. 5**). These results have major implications for structural and functional studies of the uniporter. Indeed, members of the orthologous MCU complex in basal fungi constitute relevant targets for future research and comparative structural analyses, particularly for identifying key MCU-EMRE interactions. Our results show that human MCU can function in the presence of both human and fungal EMRE, whereas animal-like MCUs such as Am-MCU and Sp-MCU can only reconstitute mt-Ca^2+^ uptake when co-expressed with their corresponding fungal EMRE. These findings indicate a tight co-evolution between MCU and EMRE proteins, which we know to functionally and physically interact, and provides the framework to understand the sequence determinants of this interaction. Finally, our work underscores that accurate phylogenomic analyses can resolve apparent evolutionary riddles while making explicit functional predictions that can drive future experiments.

**Figure 5:**
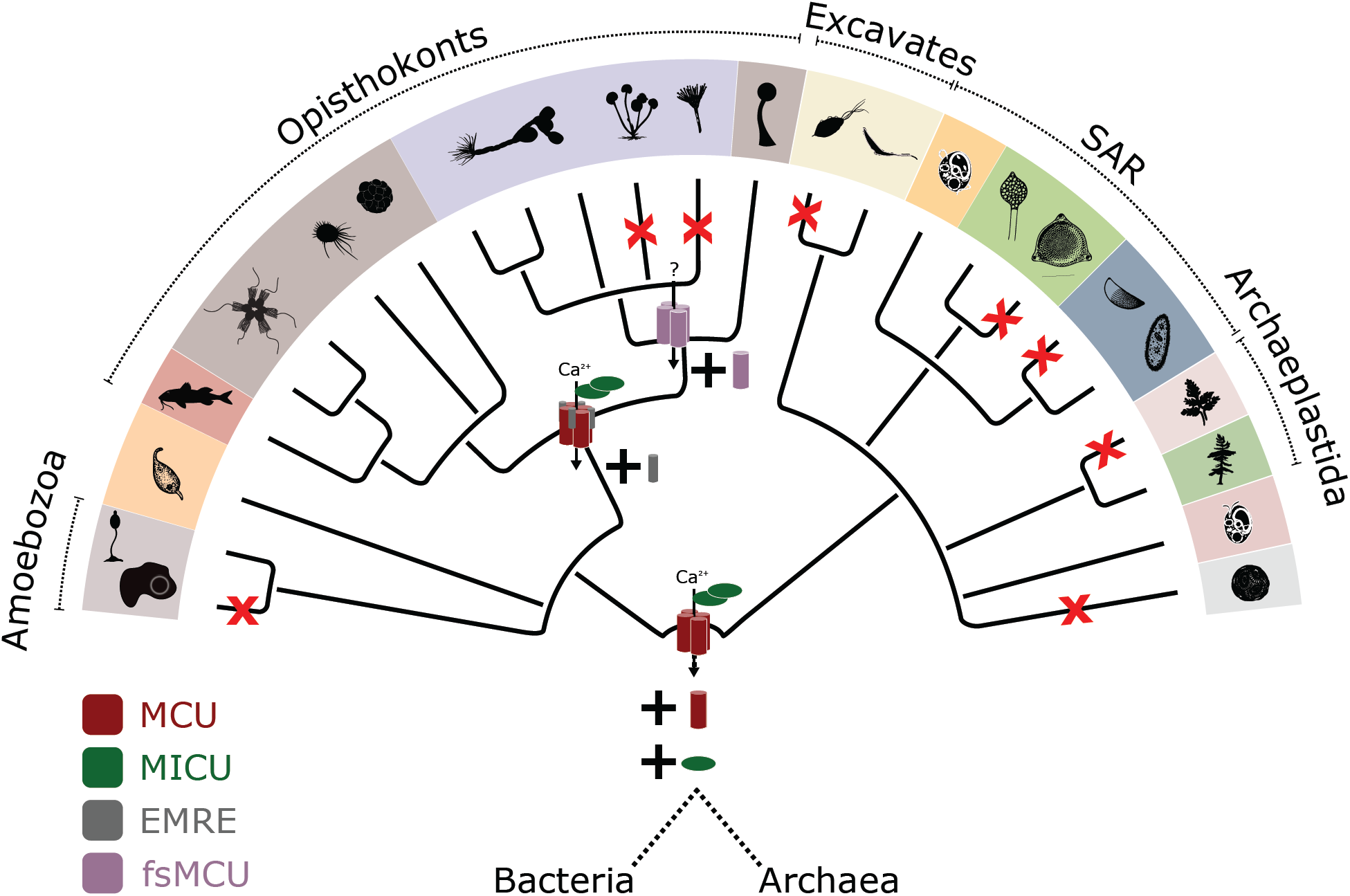
Schematic representation of the evolution of the MCU complex in Eukaryotes. Changes in the composition of the uniporter complex are shown in the respective taxonomic levels (same tree structure as in Fig. 3a). Losses of MCU in the various eukaryotic lineages are indicated in red. The color code and symbols of the different taxonomic groups are the same as in Fig. 1 and Fig. 3a, respectively.

## Methods

### Sequence data and Homology searches

The protein sequences encoded in 1,156 completely sequenced eukaryotic genomes were retrieved from Ensembl DB v91, and v37 of Ensembl Metazoa, Plants, Fungi, and Protists (see **Supplementary Table 1**). Only the genome of Catenaria anguillulae PL171 was added from Ensembl Fungi v41. For each of the protein families studied, homologs were selected on the basis of sequence similarity and phylogenetic analysis. HMMER searches were performed using HMMER 3.1b2^35^ and using the Gathering Cut-Off threshold (--cut_ga) when the raw HMM profile from Pfam was used, an e-value threshold of 10^−2^ otherwise. For all BLAST searches low complexity regions in the query sequence (default parameter) were filtered out to minimise the number of false positives and an E-value threshold of 10^−5^ was used. Conserved domains in all retrieved sequences, were annotated using the HMM profiles of Pfam release 30.0.

#### MCU

Proteins in our database containing at least one MCU (Pfam:PF04678) domain were detected using an HMMsearch. 1,076 protein sequences were selected for subsequent analysis.

#### MICU

2,105 protein sequences were retrieved from a BLAST search using Hs-MICU1 (Uniprot: Q9BPX6) as a query. HMMscan was used to search for additional domains in the retrieved sequences using all the Pfam domain profiles, and all the sequences with detected at least one Mito_carr domain (Pfam:PF00153) were classified as members of the mitochondrial carrier family (slc25a12-Aralar homologs). The Aralar-related sequences (Mito_carr domain containing) were clustered together clearly as a monophyletic clade in a phylogenetic tree, in the exclusion of known MICU sequences, and were excluded. The remaining 651 MICU sequences were re-aligned and a new phylogenetic tree with the same methods was reconstructed.

#### EMRE

EMRE sequences are characterized by a DDDD (Pfam:PF10161) domain. Their short length and low sequence conservation makes detection strikingly difficult, which explains why in some cases EMRE appears to be missing even from some animal genomes. HMMER searches with the “gathering” (--cut_ga) threshold were performed, and all detected homologs were retrieved and re-aligned, for a new HMM profile to be built and used to search back the genome database in a second iteration. One EMRE sequence was detected in *A.macrogynus* in the first search, while one more sequence in the same species, as well as in *S.punctatus* and *C.anguillulae*, were detected in the second iteration. The detected EMRE sequences in this second iteration are those that were further considered for all analyses.

#### NCLX (NCKX6)

Na_Ca_ex Pfam domain characterizes an ubiquitous superfamily of sodium/calcium exchangers that regulate intracellular Ca^2+^ concentrations in many cell types. Therefore selecting on that domain of the NCKX6 (NCLX) homologs using HMMER returned 4024 hits. To narrow down the number of hits for more accurate alignment and phylogenetic reconstruction, we used the human NCLX sequence as a query (uniprot:Q6J4K2) for a blast search, retrieving 2,105 sequences for phylogenetic analysis. Using the human members as reference, 1,391 sequences across eukaryotes were selected as related to the NCLX clade (NCLX orthologs).

### Phylogenetic analysis

Sequences of all the protein families described were aligned, and the alignment was trimmed and used to compute a phylogenetic tree. The selected homologous proteins were aligned with MAFFT v7.394^36^ (E-INS-I for MICU, EMRE, and NCLX families and L-INS-i for the MCU family, based on their multi/single-domain architecture) and a soft trimming was applied, filtering out positions in the alignment with gaps in more than 99% of the sequences using trimAl^37^ (-gt 0.01). IQ-TREE v1.6.8^38^ was used to derive Maximum Likelihood (ML) trees. LG was selected as the base model to test all rate-heterogeneous models using the “-mset LG” parameter (26 protein models, combinations of invariable sites +I, discrete Gamma model with 4 rate categories +G, the FreeRate model +R with 2-10 categories, and empirical AA frequencies estimated by the data). The best-fit models, chosen according to the Bayesian information criterion (BIC), were LG+R10 for the MCU and MICU alignments, LG+F+R10 for NCLX alignment. Branch supports were obtained using the ultrafast bootstrap implemented in the IQ-TREE program with 1000 replicates. The ETE Toolkit^33^ was used for all taxonomic and phylogenetic tree operations and visualization. Sequence alignments were visualized using Jalview2^39^, and the Multi-Harmony^34^ method was used identify patterns of variation across the different protein clades, positions conserved across the animal MCUs and the animal-like fungal MCUs but not in the fsMCUs.

### Cell lines

HeLa cells stably expressing a wild-type mitochondrial matrix-targeted GFP-aequorin (mt-AEQ) were generated as previously described^40^. Mt-AEQ HeLa cells stably expressing Hs-MCU, Sp-fMCU, Sp-MCU, Am-fMCU1, Am-MCUa, Am-MCUb, Af-MCU, and Nc-MCU from the pLX304 lentiviral vector on either wild-type or MCU knock-down (shMCU) mt-AEQ HeLa cells were generated as previously described^25^. cDNAs for Sp-fMCU, Sp-MCU, Am-fMCU1, Am-MCUa, and Am-MCUb expression without a stop codon were codon optimized for human expression, synthesized de novo in the PuC57 vector and amplified with flanked attB1 and attB2 sites by PCR (see **Supplementary Table 2**). PCR products were first integrated into the pDONR221 vector and then into the pLX304 destination vector by site-specific recombination according to manufacturer’s instructions (Life Technologies). All cell lines were grown in high-glucose Dulbecco’s modified Eagle’s medium (DMEM) supplemented with 10% FBS at 37°C and 5% CO_2_.

### Yeast Strains

A yeast strain expressing mt-AEQ with Hs-MCU and Hs-EMRE was generated as in^25^ and^40^. To generate yeast strains co-expressing mt-AEQ together with different combinations of fungal and human MCU and EMRE orthologs, cDNAs were amplified from the pLX304 vector (see **Supplementary Table 3**) and cloned into the yeast expression plasmids p423GPD (Hs-MCU, Sp-fsMCU, Sp-MCU, Am-fsMCU1, Am-MCUa, Am-MCUb, Ce-MCU, Fg-MCU, Ma-MCU, and Nf-MCU) and p425GPD (Hs-EMRE, wild-type and C-terminal domain truncated Sp-EMRE, Am-EMRE1 and Am-EMRE2, and Hs-EMRE with an addition of the extra C-terminal domain from Sp-EMRE). The YPH499 strain was then transformed with the respective plasmids and transformants were selected on synthetic dextrose plates, with adenine, lysine and tryptophan as selection markers.

### MCU and EMRE knockout in HeLa

mt-AEQ HeLa cells with knockout of MCU were generated by CRISPR targeting^41^. In brief, a cDNA encoding NLS-Cas9 was isolated from pX330 (Addgene #42230) by EcoRI digestion and cloned upstream of the ubiquitous CAG promoter. Two sets of single-guide RNA (sgRNA) were designed using an online tool CRISPOR (MCU-KO1 gRNAa: 5’-CAGGAGCGATCTACCTGCGG-3’; MCU-KO2 gRNAa: 5’-TGAACTGACAGCGTTCACGC-3’) and cloned into a pX330-Puro-ccdB vector as previously described^42^. After which, pX330-Puro-ccdB vector containing sgRNAs were transfected into mt-AEQ HeLa cells using the Lipofectamine 3000 Reagent (Life Technologies), according to the manufacturer’s instructions and selected with 2 µg/mL puromycin for two days. Cells were then seeded at a density of 1 cell per well and expanded. Gene knockout was screened by PCR (MCU-KO Forward1: 5’-GCGTGTAGTTGAGAGTTACAGC-3’;MCU-KO Forward2:5’-TTTTATAAGCCAGTTCCCAGAATAACCT-3’; MCU-KO Reverse: 5’-GTTCATCCTTGCTCATGGCATT-3’) and confirmed by sequencing and western blot.

### Isolation of Crude Mitochondria from HeLa Cells

Crude mitochondria were isolated from HeLa cells as previously described^25^. Briefly, HeLa cells were grown to confluency, rinsed with PBS and resuspended in ice-cold isolation buffer (IB: 220 mM mannitol, 70 mM sucrose, 5 mM HEPES-KOH pH 7.4, 1 mM EGTA-KOH pH 7.4, protease inhibitors). Cells were permeabilized by nitrogen cavitation at 600 psi for 10 minutes at 4°C and then centrifuged at 600 x g for 10 minutes. The supernatant was transferred into new tubes and centrifuged at 8000 x g for 10 minutes at 4 °C. The resulting pellet containing crude mitochondria was resuspended in IB for protein topology analysis.

### Analysis of Mitochondrial Protein Topology

Proteinase K (PK) protection assay was performed on mitochondria isolated from HeLa cells as previously described^25^. Roughly, 30 µg of freshly isolated mitochondria were gently resuspended in 30 µl of IB buffer with either increasing concentrations of digitonin or 1% Triton X-100 in the presence of 100 µg/ml PK and incubated at room temperature for 15 minutes. The reaction was stopped by the addition of 5 mM PMSF, followed by incubation on ice for 10 minutes. Samples were mixed with 10 µl of 4 X Laemmli buffer containing 10 % 2-mercaptoethanol and boiled for 5 minutes at 98 °C for immunoblot analysis.

### Subcellular Fractionation of Yeast Cells

Expression and subcellular localization of heterologous expressed proteins in yeast was tested by immunoblot analysis of cytosolic and mitochondrial fractions isolated from recombinant yeast strains as previously described^25^. Briefly, yeast cells were grown at 30°C in a selective lactate medium supplemented with the respective selection markers till an OD ∼0.8. The cell pellet was re-suspended in a buffer containing buffer 0.6 M sorbitol, 20 mM HEPES/KOH pH 7.2, 80 mM KCl, and 1 mM PMSF, and vortexed five times for 30 seconds with glass beads (425-600 µm diameter), with a 30 seconds cooling interval in between to break cell wall and plasma membrane. After a first centrifugation step at 1000 *g* for 5 minutes at 4°C, the supernatant was further centrifuged at 20,000 *g* for 10 minutes at 4°C to obtain the mitochondrial fraction (pellet). The supernatant (cytosolic fraction) was precipitated with trichloroacetic acid at -20°C for 1 hour, washed once with cold acetone and centrifuged at 20,000 g for 10 minutes at 4°C.

### Measurements of Mitochondrial Calcium Uptake in Yeast and HeLa Cells

*In vivo* analyses of mitochondrial Ca^2+^ uptake in intact yeast cells were performed as previously described^25^. Briefly, yeast cells were collected at an OD ∼0.8, washed three times with milliQ water and starved for 1.5 hours at room temperature in a nutrient-free buffer (NFB, 100 mM Tris, pH 6.5 (1×10^8^ cells/mL). Afterwards, cells were collected at 3,500 rpm for 5 minutes and resuspended in NFB to a higher density (25×10^8^ cells/mL) in the presence of 50 µM native coelenterazine (Abcam, ab145165) to reconstitute the photoprotein aequorin. After 30 minutes in the dark at room temperature, 0.5×10^8^ cells/well were plated into a white 96-well plate and Ca^2+^-dependent light kinetics was recorded upon stimulation with 1 mM CaCl_2_ and 100 mM glucose, at 0.5 seconds interval in a MicroBeta2 LumiJET Microplate Counter. At the end of each experiment, cells were lysed with 1 mM digitonin for 5 minutes at 37°C and any residual aequorin counts were collected upon the addition of CaCl_2_ to a final concentration of 140 mM. Mitochondrial Ca^2+^ uptake was measured in mt-AEQ HeLa cells as previously described^40^.

### Quantification of Calcium Transients

Quantification of mt-Ca^2+^ concentration was performed using a MATLAB software as previously described in ^40^. The dynamics of mt-Ca^2+^-dependent luminescence signal was smoothed by the cubic spline function:

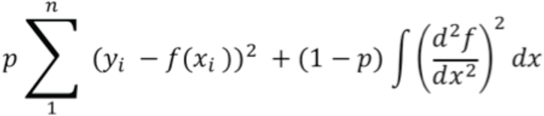

Where, p is a smoothing parameter, controlling the tradeoff between fidelity to the data and roughness of the function estimate, f is the estimated cubic spline function to minimize the above function, and x_i_ and y_i_ are the dynamical data points. Here, p is set at 0.5. Parametrization of the Ca^2+^-dependent luminescence kinetics was performed in order to determine the maximal amplitude of the luminescence signal (peak) and the left slope of the bell-shaped kinetic trace. Aequorin-based luminescence signal calibration into mt-Ca^2+^ concentration was performed using the algorithm reported in ^43^ for wild-type aequorin and native coelenterazine, with the following formula:

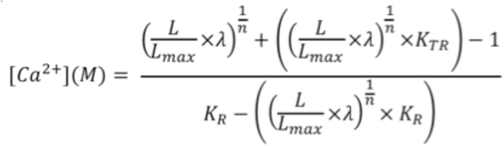

Where l= 1, K_R_= 7.23×10^6^, K_TR_= 120 and n = 2.99 are the calibration values used for WT aequorin and native coelenterazine.

### Data Analysis

Data are represented as mean ± SEM and the statistical analysis of each experiment is described in the figure legends including the statistical tests used and the exact value of biological replicates. For each biological replicate experiment at least 3 technical replicates were used for quantification and data analysis. Normal distribution was tested by Shapiro-Wilk normality test. Statistical tests between multiple datasets and conditions were carried out using one-way analysis of variance (ANOVA) followed by Dunnett’s Multiple Comparison tests. Statistical analyses were performed using GraphPad Prism (GraphPad Software, version 7).

## Acknowledgements

TG group acknowledges support from the Spanish Ministry of Economy, Industry, and Competitiveness (MEIC) for the EMBL partnership, and grants ‘Centro de Excelencia Severo Ochoa 2013-2017’ SEV-2012-0208, and BFU2015-67107 co-founded by European Regional Development Fund (ERDF); from the CERCA Programme / Generalitat de Catalunya; from the Catalan Research Agency (AGAUR) SGR857, and grants from the European Union’s Horizon 2020 research and innovation programme under the grant agreement ERC-2016-724173. TG also receives support from an INB Grant (PT17/0009/0023 – ISCIII-SGEFI/ERDF). FP group was supported by the Munich Center for Systems Neurology (SyNergy EXC 2145 /ID 390857198) and ExNet-0041-Phase2-3 ("SyNergy-HMGU”)’ through the Initiative and Network Fund of the Helmholtz Association to F.P.; The Bert L & N Kuggie Vallee Foundation (to F.P. and J.W.); the Juniorverbund in der Systemmedizin ‘mitOmics’ (FKZ 01ZX1405B to V.G.). A.A.P. was supported by a postdoctoral research fellowship from EMBO (118-2017) while writing this article. A.C.S. was partially supported by the Aging and Metabolic Programming project (AMPro).

## Author contributions

A.A.P., T.G., and F.P. conceived the project, designed the experiments and wrote the manuscript. V.G. performed most of the experiments with help from J.W. and A.C.S and A.A.P. performed computational analysis. T.G. and F.P. supervised work and acquired funding.

---

**Extended Data Figure 1.**
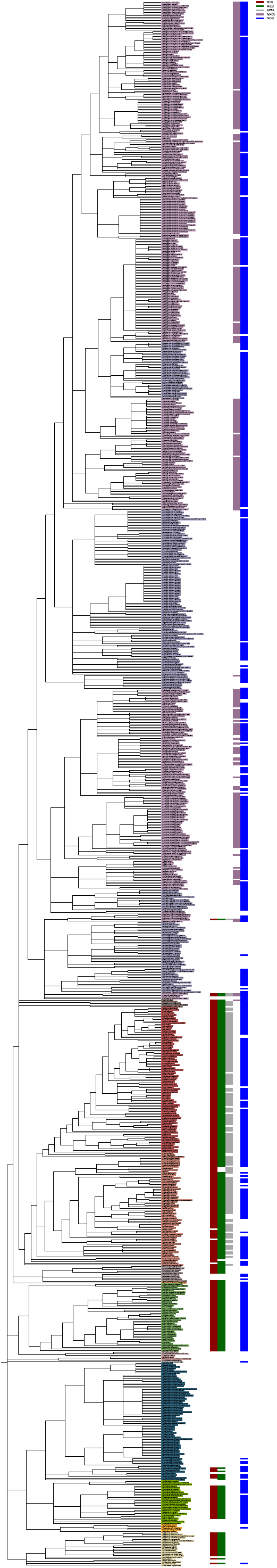
Phylogenetic distribution of the mitochondrial calcium transporter complex protein families. The tree is collapsed to the species level, resulting in 969 species. Extended version of Figure 1, including also the distribution of NCLX, which appears to be largely uncoupled to that of the MCU complex members.

**Extended Data Figure 2.**
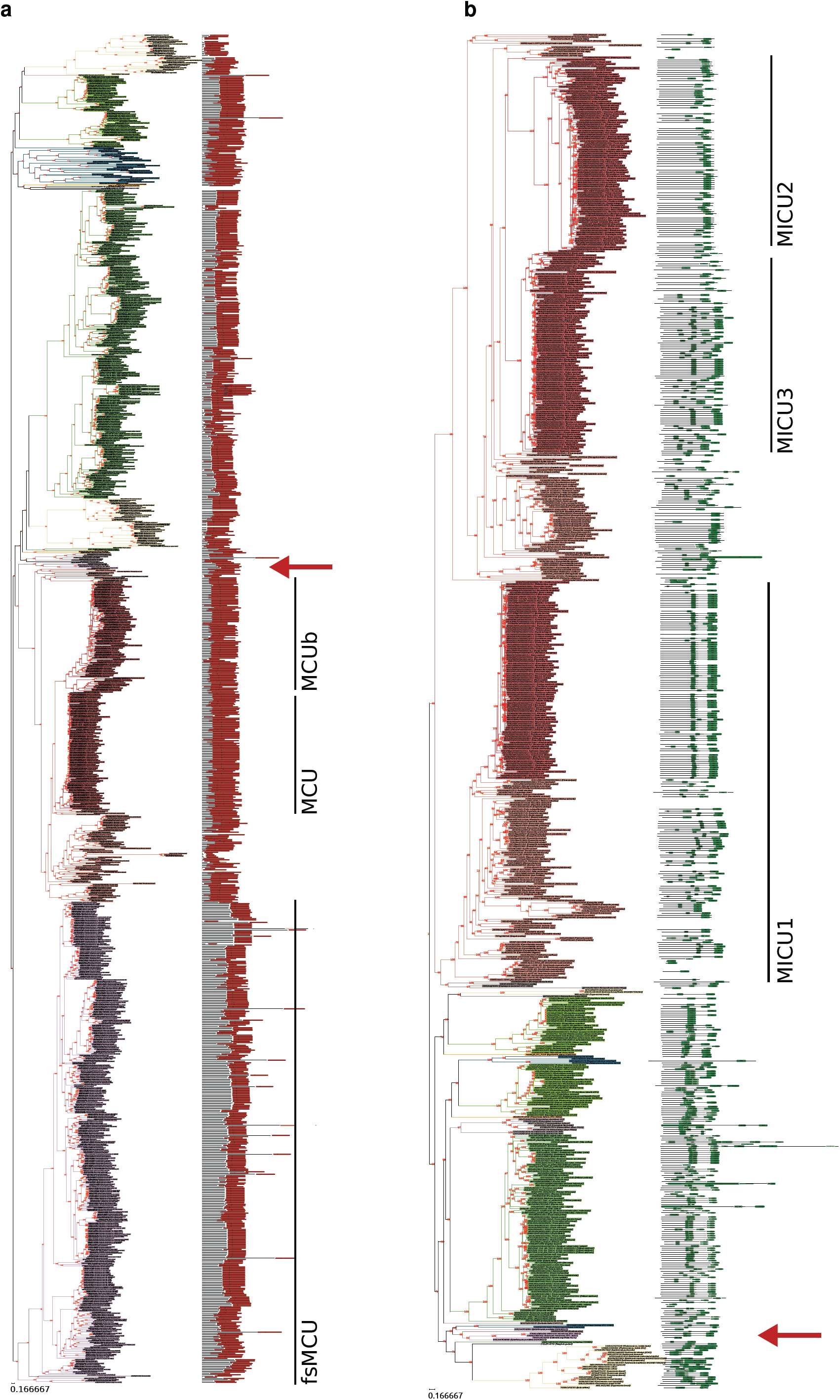
Maximum likelihood (ML) phylogenetic trees of MCU and MICU families. Phylogenies of the 1,064 MCU sequences (**a**) and the 651 MICU sequences (**b**). Full version with sequences ids of Fig. 2a,b. The UFBoot (Ultrafast Bootstrap Approximation) support values, as implemented in IQ-TREE 1.6.8, are indicated in red. The main subfamilies are shown, based on the human representatives. The basic domain architecture according to Pfam is plotted on the right. The MCU family is characterized by the presence of one “MCU” domain (in red), while the typical MICU sequence consists of two “EF-hand” domains (in green). In both (**a**) and (**b**), the position of the animal related MCU and MICU sequences from fungi and *F.alba* are indicated with a red arrow. See also Methods.

**Extended Data Fig. 3:**
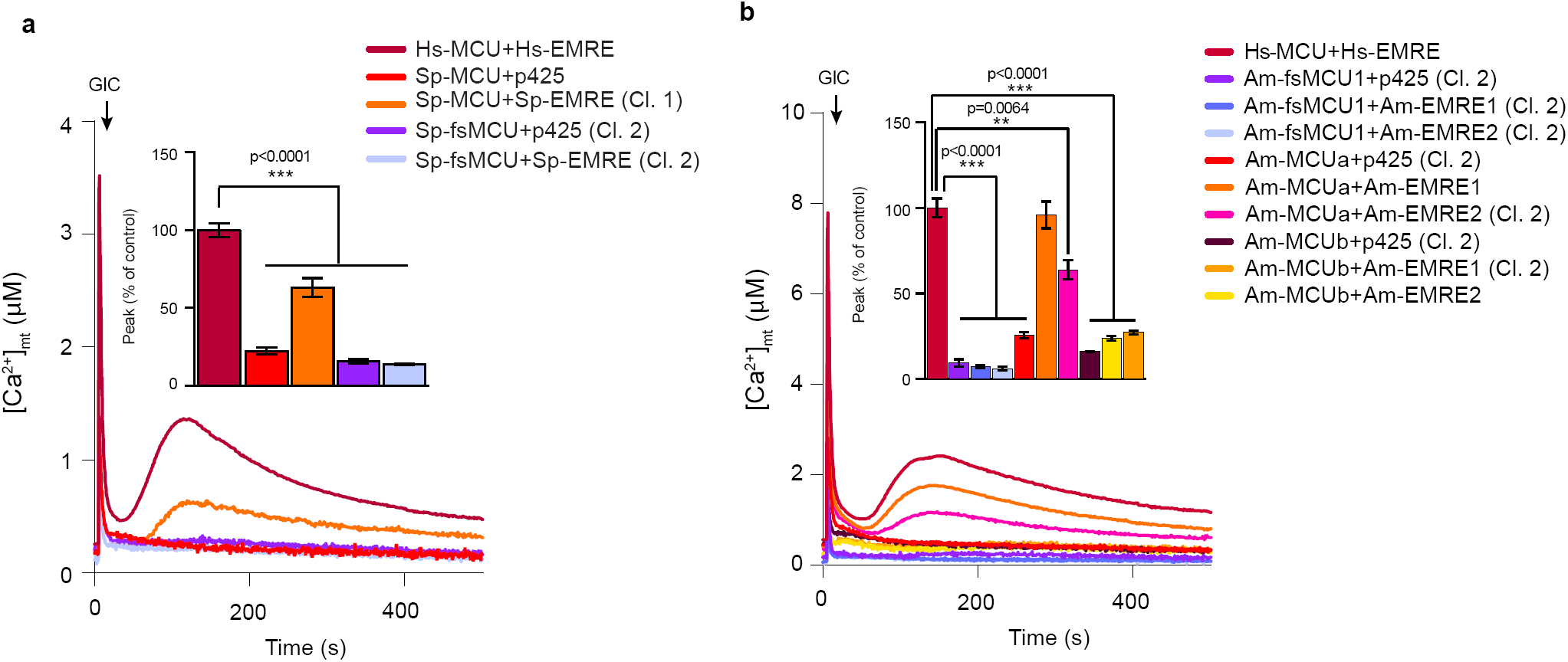
Reconstitution of mt-Ca^2+^ uptake in yeast cells expressing *S. punctatus* or *A. macrogynus* MCU and EMRE orthologs. **a,b**, Representative traces and quantification of mt-Ca^2+^ transients in yeast cells expressing animal-like or fungal-specific MCU orthologs from *S. punctatus* (**a**) and *A. macrogynus* (**b**) with either their respective EMREs or empty vector (p425) upon glucose-induced calcium (GIC) stimulation in presence of 1 mM CaCl_2_. All data represent mean ± SEM (n=3); ***p < 0.0001, **p < 0.001, one-way ANOVA with Dunnett’s Multiple Comparisons Test.

**Extended Data Fig. 4.**
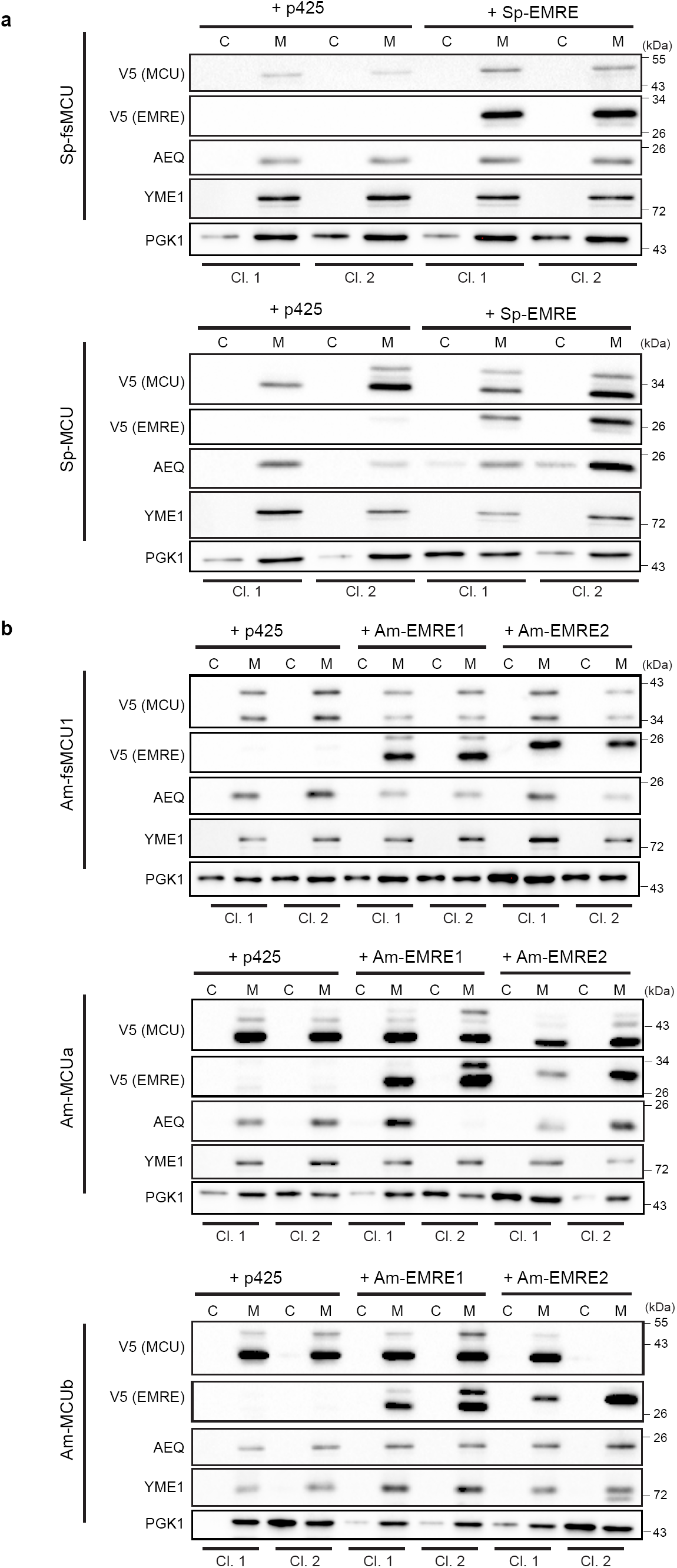
Heterologous expression of *S. punctatus* and *A. macrogynus* MCU and EMRE orthologs in yeast. **a,b**, Immunoblot analysis of cytosolic (C) and mitochondrial (M) fractions isolated from yeast clones (Cl.) expressing mt-AEQ together with either (**a**) *S. punctatus* MCU (Sp-fsMCU, Sp-MCU) and EMRE (Sp-EMRE) orthologs or (**b**) *A. macrogynus* MCU (Am-fsMCU1, Am-MCUa, Am-MCUb) and EMRE (Am-EMRE1, Am-EMRE2) orthologs fused to a C-terminal V5-tag, using the following antibodies: α-V5 (Life Technologies, R96025), α-AEQ (Merck/Millipore, MAB4405), α-YME1, PGK1 (Life Technologies, 459250). YME1 was used as control for yeast mitochondrial targeted protein and PGK1 was used as control for cytosolic protein.

**Extended Data Fig. 5.**
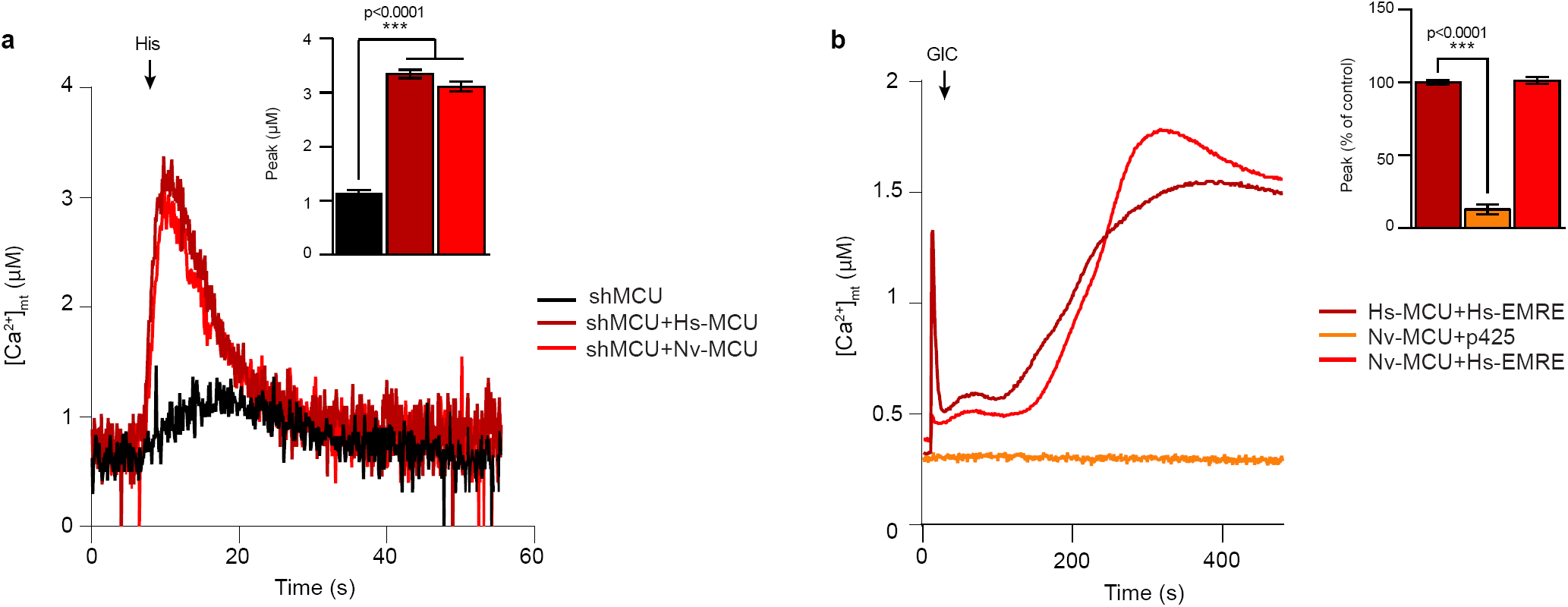
Reconstitution of mt-Ca^2+^ uptake in yeast and HeLa cells expressing *N. vectensis* MCU. **a**, Representative traces and quantification of mt-Ca^2+^ transients in MCU knockdown (shMCU) HeLa cells expressing *N. vectensis* MCU upon histamine (His) stimulation. **b**, Representative traces and quantification of mt-Ca^2+^ transients in yeast cells expressing *N. vectensis* MCU with either empty vector (p425) or human EMRE upon glucose-induced calcium (GIC) stimulation in presence of 1 mM CaCl_2_. All data represent mean ± SEM; n=3-6; ***p < 0.0001, one-way ANOVA with Dunnett’s Multiple Comparisons Test.

**Extended Data Figure 6.**
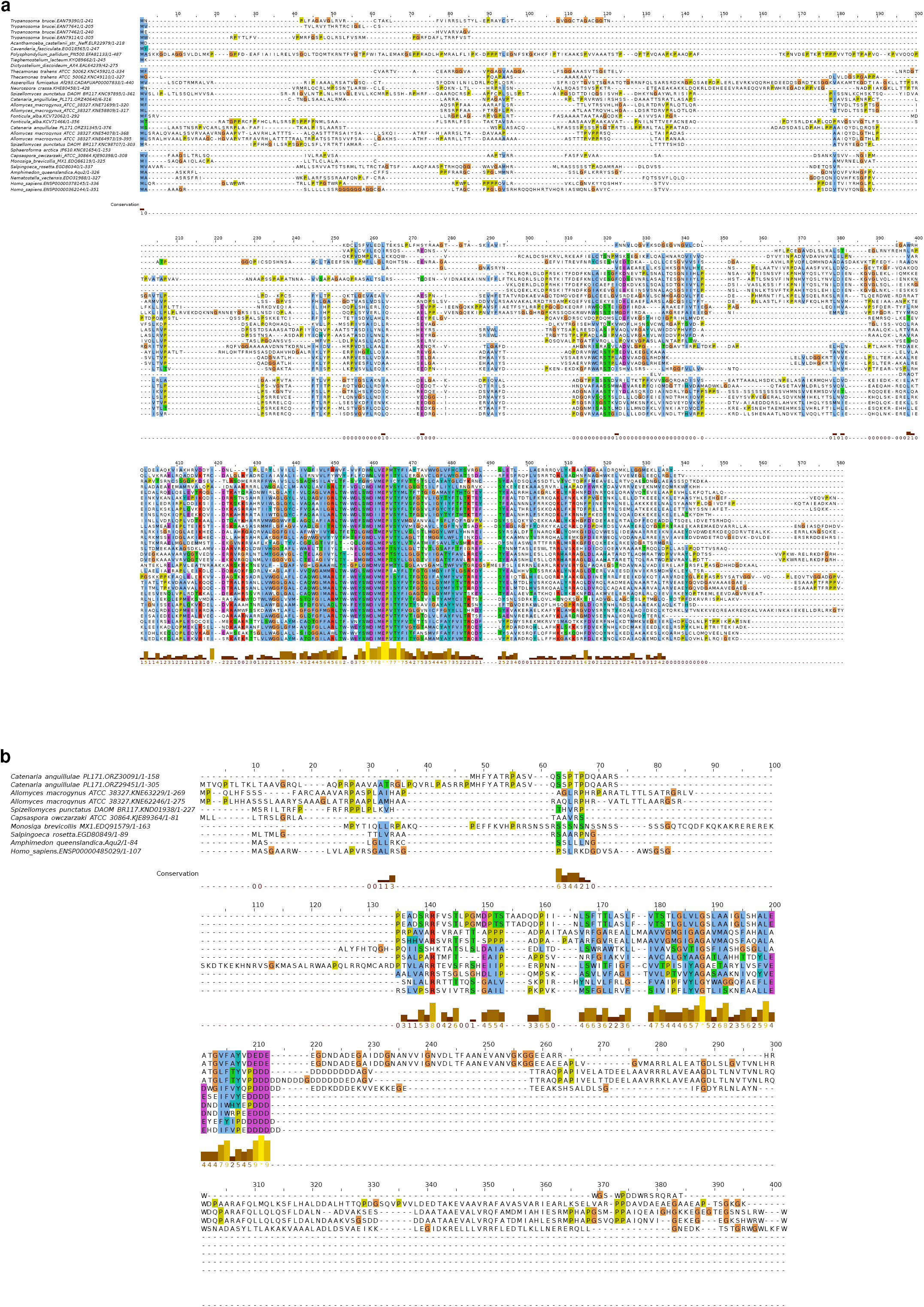
**a,b**, Multiple sequence alignments of opisthokont members of MCU (**a**) and EMRE (**b**) families. In (**a**) MCU regions poorly aligned or specific to non-opisthokont species have been removed, whereas in (**b**) the full EMRE sequences are shown, for all the species in (**a**) where EMRE was detected.

**Extended Data Fig. 7.**
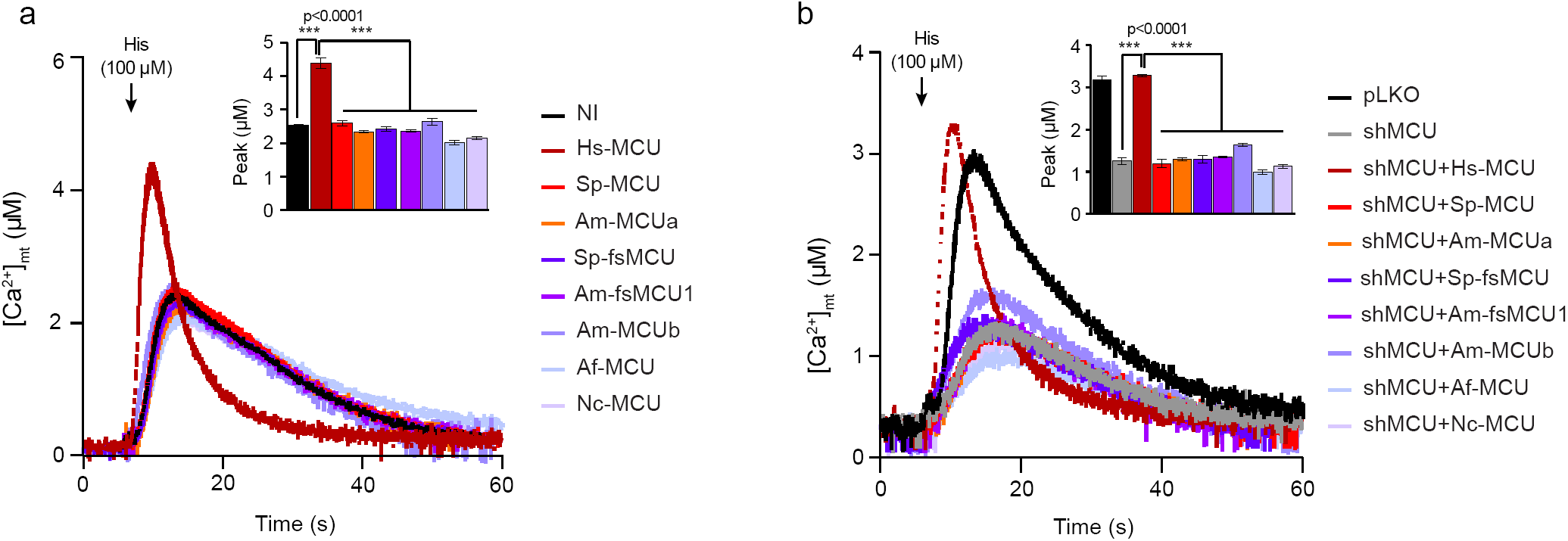
Reconstitution of mt-Ca^2+^ uptake in HeLa cells expressing fungal MCU orthologs. **a,b**, Quantification of mt-Ca^2+^ transients in either wild-type (a) or MCU knockdown (shMCU) (b) HeLa cells expressing different fungal MCU orthologs upon histamine (His) stimulation. All data represent mean ± SEM; n=4; ***p < 0.0001, one-way ANOVA with Dunnett’s Multiple Comparisons Test.

**Extended Data Fig. 8.**
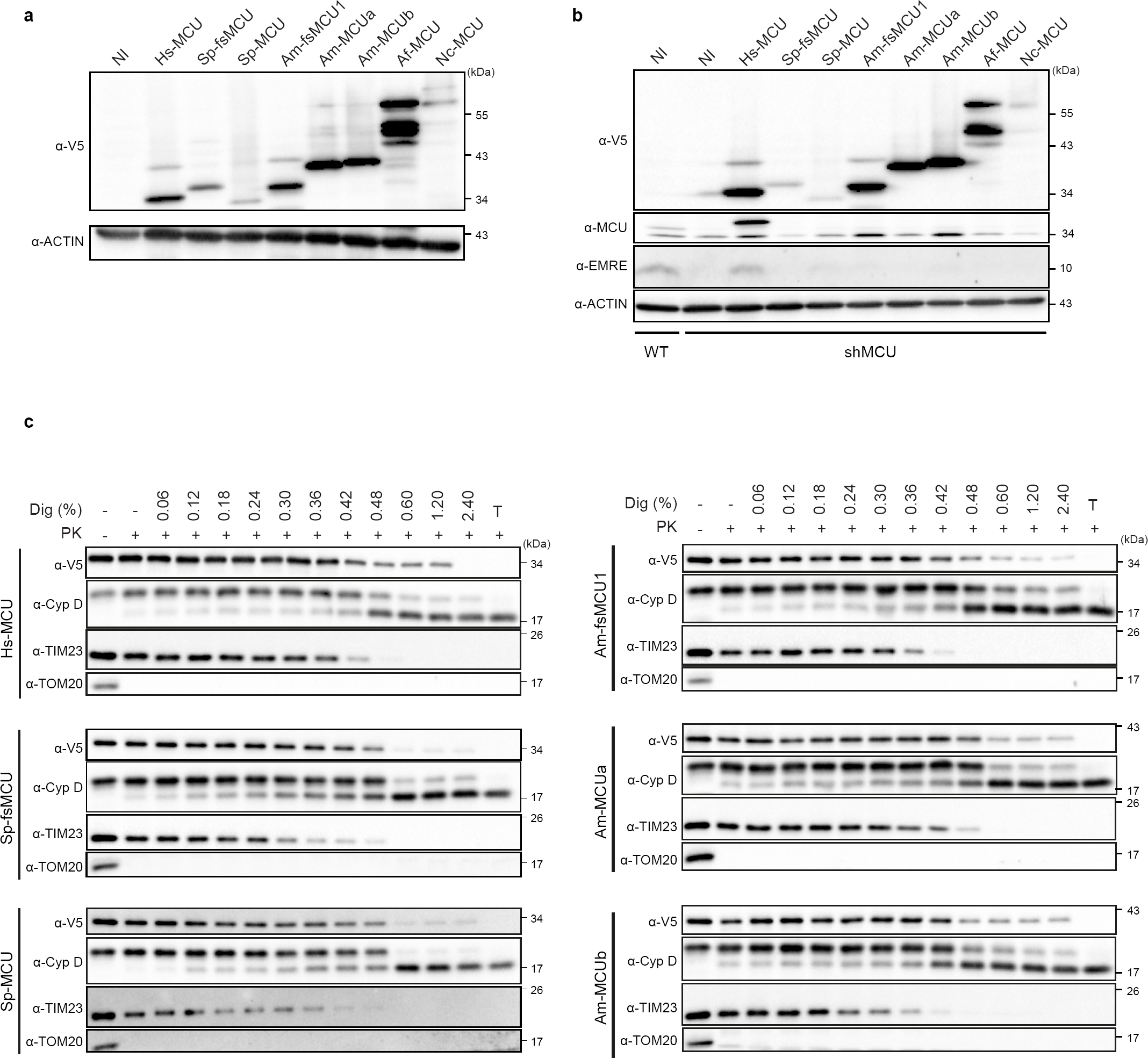
Expression and localization of fungal MCU orthologs in HeLa cells. Immunoblot analysis of whole cell lysates from wild-type (**a**) and MCU knock-down (shMCU) (**b**) HeLa mt-AEQ cells stably expressing human or fungal MCU proteins fused to a C-terminal V5 tag using the following antibodies: α-MCU (Sigma Aldrich, HPA01648), α-V5 (Life Technologies, R96025), α-EMRE (Santa Cruz Biotechnology, sc-86337), α-ACTIN (Sigma-Aldrich, A2228). NI, not infected. **c**, Analysis of protein topology by proteinase K (PK) treatment of mitochondria isolated from sh-MCU HeLa mt-AEQ cells expressing human and fungal MCU, using the following antibodies: α-V5 (Life Technologies, R96025), α-TIM23 (BD Bioscience, 611222), α-TOM20 (Abcam, ab56783), and α-Cyclophilin D (Abcam, ab110324). TOM20, TIM23, and CypD were used as controls for integral mitochondrial outer membrane, inner membrane and soluble matrix targeted proteins, respectively. T, triton (1%); Dig., digitonin.

**Extended Data Fig. 9.**
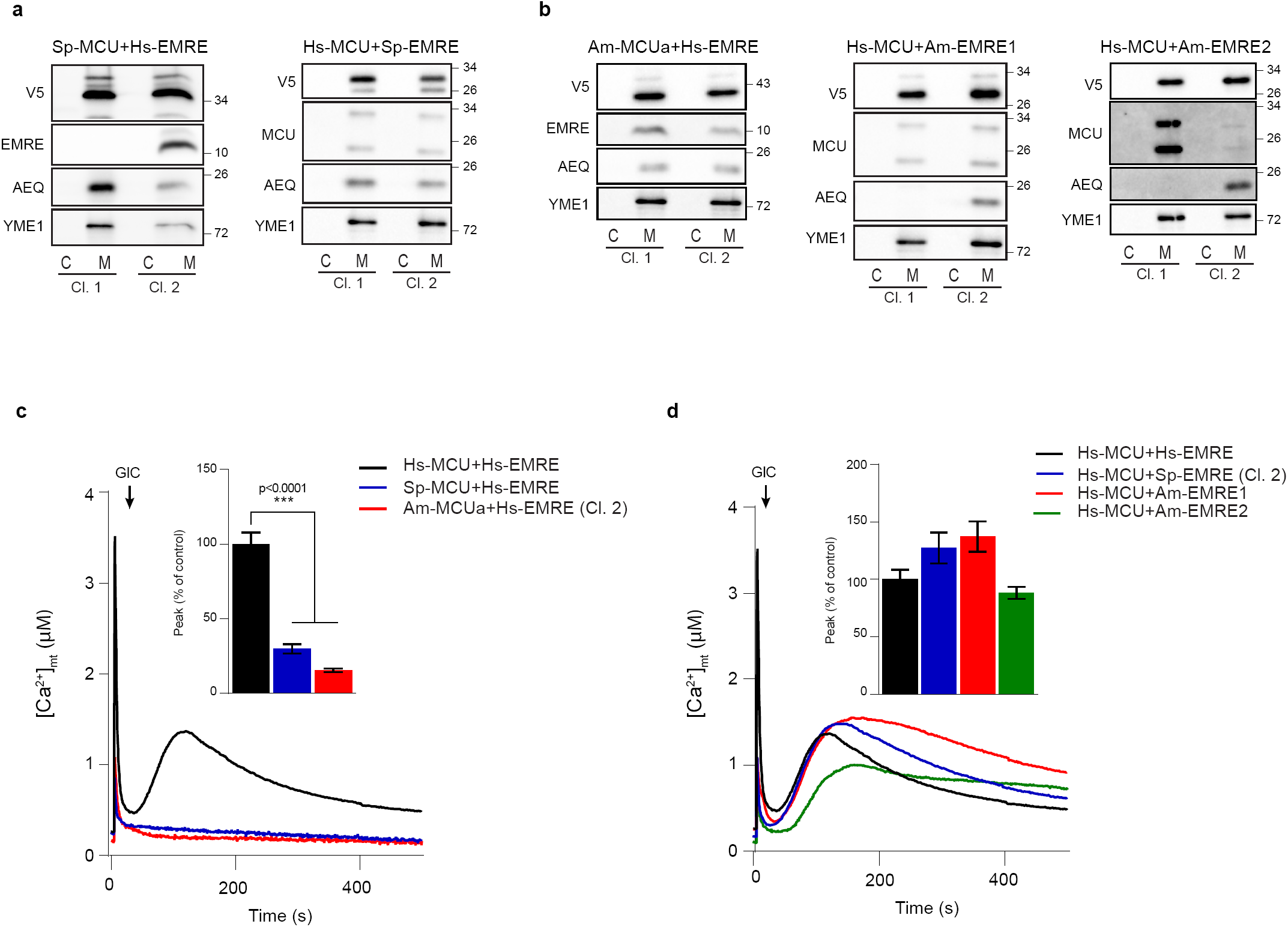
Reconstitution of mt-Ca^2+^ uptake in yeast cells expressing human MCU and EMRE with either fungal EMRE or animal-like MCU orthologs from S. punctatus and A. macrogynus respectively. **a,b**, Immunoblot analysis of cytosolic (C) and mitochondrial (M) fractions isolated from yeast clones (Cl.) expressing mt-AEQ together with either (**a**) S. punctatus or (**b**) A. macrogynus MCU and EMRE orthologs fused to a C-terminal V5-tag. YME1 was used as control for yeast mitochondrial targeted protein. **c,d**, Representative traces and quantification of mt-Ca^2+^ transients in yeast cells expressing either human and animal-like S. punctatus and A. macrogynus MCU with human EMRE (**c**) or species-specific EMRE with human MCU (**d**) upon glucose-induced calcium (GIC) stimulation in presence of 1 mM CaCl2. All data represent mean ± SEM (n=3); ***p < 0.0001, one-way ANOVA with Dunnett’s Multiple Comparisons Test.

**Extended Data Fig. 10.**
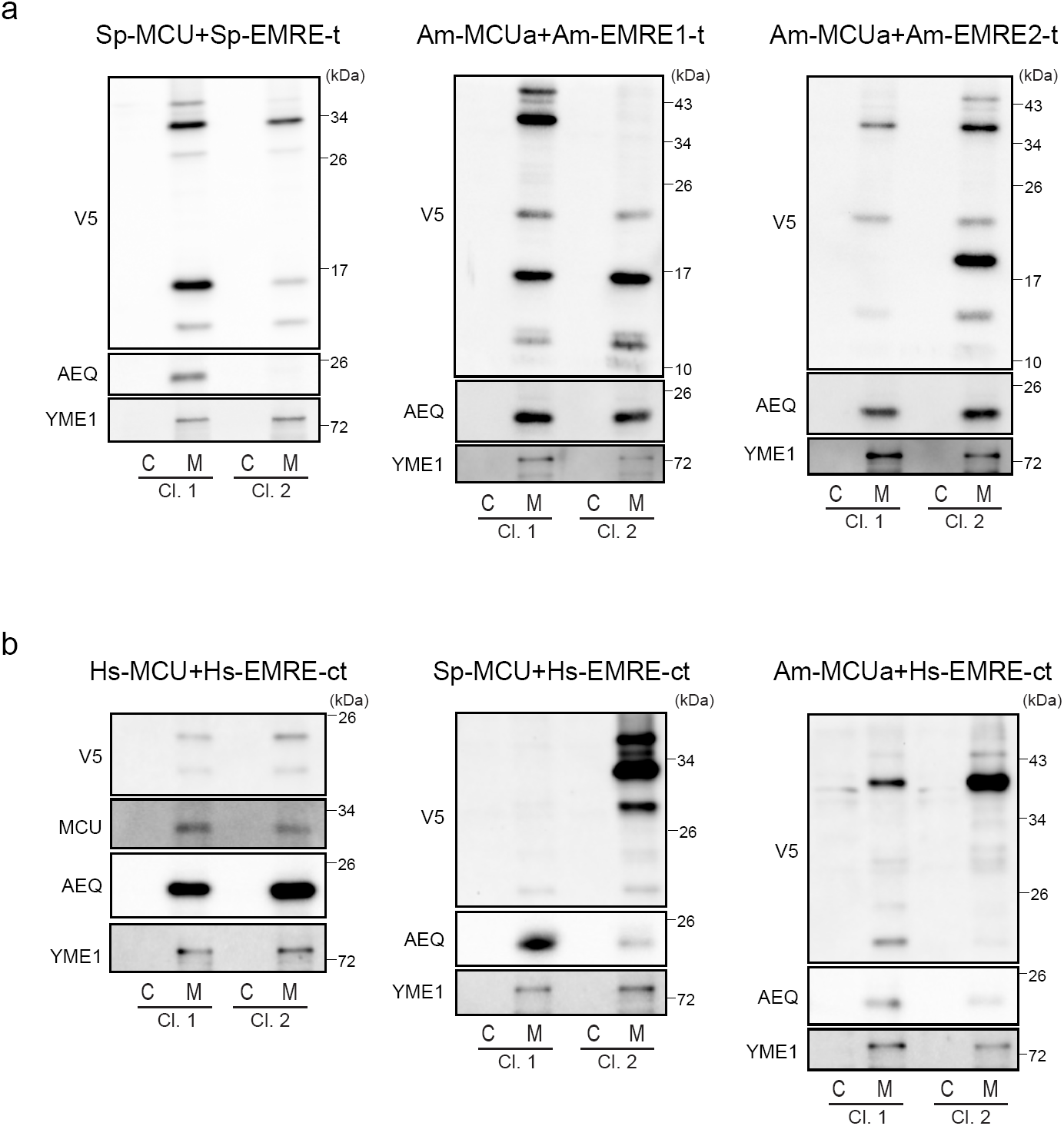
Heterologous expression of *S. punctatus* and *A. macrogynus* MCU and EMRE orthologs in yeast. **a,b**, Immunoblot analysis of cytosolic (C) and mitochondrial (M) fractions isolated from yeast clones (Cl.) expressing mt-AEQ together with either (**a**) *S. punctatus* (Sp-MCU) and *A. macrogynus* (Am-MCUa) MCU and their respective truncated EMRE (Sp-EMRE-t, Am-EMRE1-t, Am-EMRE2-t) or (**b**) Human, *S. punctatus* (Sp-MCU) *and A. macrogynus* (Am-MCUa) MCU and human EMRE with an added fungal extra C-terminal domain (Hs-EMRE-ct), fused to a C-terminal V5-tag, using the following antibodies: α-V5 (Life Technologies, R96025), α-AEQ (Merck/Millipore, MAB4405), α-YME1. YME1 was used as control for yeast mitochondrial targeted protein.

**Extended Data Fig. 11.**
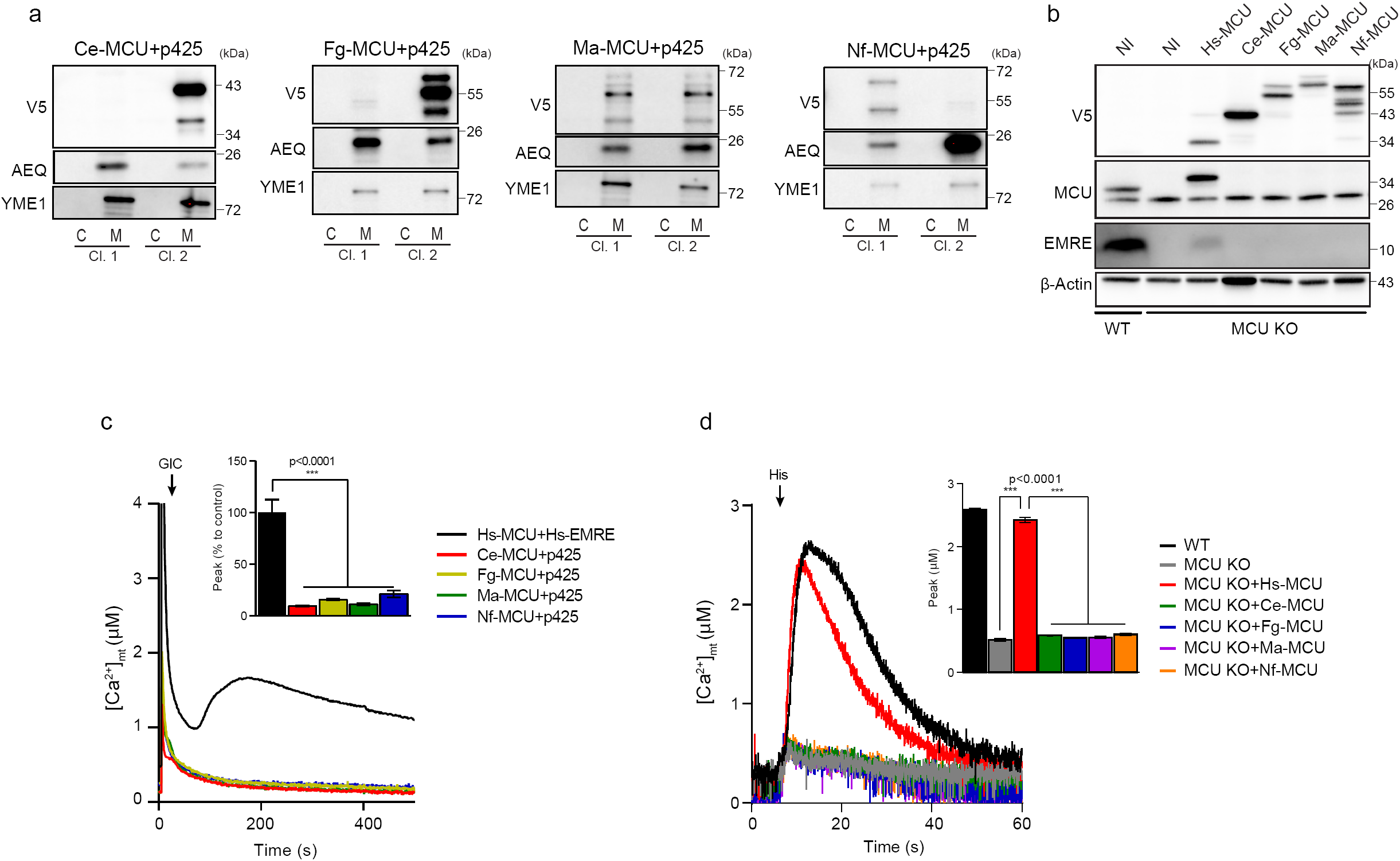
Reconstitution of mt-Ca^2+^ uptake in yeast and HeLa cells expressing fungal MCU orthologs. **a**, Immunoblot analysis of cytosolic (C) and mitochondrial (M) fractions isolated from yeast clones (Cl.) expressing mt-AEQ together with C. europaea (Ce-MCU), F. graminearum (Fg-MCU), M. acridum (Ma-MCU) or N. fischeri (Nf-MCU) MCU fused to a C-terminal V5-tag and an empty vector (p425). YME1 was used as control for yeast mitochondrial targeted protein. **b**, Immunoblot analysis of whole cell lysates from wild-type (WT) or MCU knock-out (MCU KO) HeLa mt-AEQ cells stably expressing human or fungal MCU proteins fused to a C-terminal V5 tag using the following antibodies: α-MCU (Sigma Aldrich, HPA01648), α-V5 (Life Technologies, R96025), α-EMRE (Santa Cruz Biotechnology, sc-86337), α-ACTIN (Sigma-Aldrich, A2228). NI, not infected. **c**, Representative traces and quantification of mt-Ca^2+^ transients in yeast cells expressing fungal MCU (Ce-MCU, Fg-MCU, Ma-MCU or Nf-MCU) and an empty vector (p425) upon glucose-induced calcium (GIC) stimulation in presence of 1 mM CaCl2. **d**, Quantification of mt-Ca^2+^ transients in WT or MCU KO HeLa mt-AEQ cells expressing different fungal MCU orthologs (Ce-MCU, Fg-MCU, Ma-MCU or Nf-MCU) upon histamine (His) stimulation. All data represent mean ± SEM (n=4); ***p < 0.0001, one-way ANOVA with Dunnett’s Multiple Comparisons Test.

