## Supplementary Tables 2 and 3 for "Discovery of EMRE in fungi resolves the true evolutionary history of the mitochondrial calcium uniporter"

**Supplementary Table S2**. List of PCR primers used to amplify cDNA from PuC57 vector and perform Gateway cloning into pLX304 destination vector.

| Species | cDNA | Direction | Primer sequence |
|---|---|---|---|
| *Spizellomyces punctatus* | Sp-MCU | Forward | 5'-GGG GAC AAG TTT GTA CAA AAA AGC AGG CTT AGC CAC CAT GCG AAT CCC CTT CCA CCA CGG C-3' |
| | | Reverse | 5'-GGG GAC CAC TTT GTA CAA GAA AGC TGG GTT TGT TGC CTC CAC CCT GAC CC-3' |
| | Sp-fsMCU | Forward | 5'-GGG GAC AAG TTT GTA CAA AAA AGC AGG CTT AGC CAC CAT GGT GAG CCT GAT TCC AGT GGG GG-3' |
| | | Reverse | 5'-GGG GAC CAC TTT GTA CAA GAA AGC TGG GTT AAT CAG TCT GCC CAT TCG TCC AGC-3' |
| | Sp-EMRE | Forward | 5'-GGG GAC AAG TTT GTA CAA AAA AGC AGG CTT AGC CAC CAT GTC CCG CAT TCT GAC ACG ATT CCC A-3' |
| | | Reverse | 5'-GGG GAC CAC TTT GTA CAA GAA AGC TGG GTT CCA GAA CTT CAG CCA GCC CCA-3' |
| | Sp-EMRE-t | Forward | 5'-GGG GAC AAG TTT GTA CAA AAA AGC AGG CTT AGC CAC CAT GTC CCG CAT TCT GAC ACG ATT C-3' |
| | | Reverse | 5'-GGG GAC CAC TTT GTA CAA GAA AGC TGG GTT CAC CAC CTT CTC ATC GTC ATC CTT GT-3' |
| *Allomyces macrogynus* | Am-MCUa | Forward | 5'-GGG GAC AAG TTT GTA CAA AAA AGC AGG CTT AGC CAC CAT GCT GTC AAG GGC TCT GCA GGT CG-3' |
| | | Reverse | 5'-GGG GAC CAC TTT GTA CAA GAA AGC TGG GTT CCC CTG CTT GCC CTC GCT TGT-3' |
| | Am-MCUb | Forward | 5'-GGG GAC AAG TTT GTA CAA AAA AGC AGG CTT AGC CAC CAT GCT GAT TTC TTG TCG CCT GCT GGC T-3' |
| | | Reverse | 5'-GGG GAC CAC TTT GTA CAA GAA AGC TGG GTT TGA CTG CTG CTT TCC TGC GG-3' |
| | Am-fsMCU1 | Forward | 5'-GGG GAC AAG TTT GTA CAA AAA AGC AGG CTT AGC CAC CAT GTT CGC ACA GTC CCG CCC ATT T-3' |
| | | Reverse | 5'-GGG GAC CAC TTT GTA CAA GAA AGC TGG GTT TTT GCC GCC CAG AAT CTC GTG-3' |
| | Am-EMRE1 | Forward | 5'-GGG GAC AAG TTT GTA CAA AAA AGC AGG CTT AGC CAC CAT GCC TCA GCT GCA TTT CTC ATC ATC TTT CG-3' |
| | | Reverse | 5'-GGG GAC CAC TTT GTA CAA GAA AGC TGG GTT CCA CCA TCG CAG GCT ATT TGA CCC-3' |
| | Am-EMRE2 | Forward | 5'-GGG GAC AAG TTT GTA CAA AAA AGC AGG CTT AGC CAC CAT GCC CCC TCT GCA CCA CGC C-3' |
| | | Reverse | 5'-GGG GAC CAC TTT GTA CAA GAA AGC TGG GTT CCA CCA CCT CCA ATG GCT TTT CC-3' |
| | Am-EMRE1-t | Forward | 5'-GGG GAC AAG TTT GTA CAA AAA AGC AGG CTT AGC CAC CAT GCC TCA GCT GCA TTT CTC ATC ATC-3' |
| | | Reverse | 5'-GGG GAC CAC TTT GTA CAA GAA AGC TGG GTT CAC TCC TGC GTC ATC GTC ATC GT-3' |

| | | | |
|---|---|---|---|
| | Am-EMRE1-t | Forward | 5'-GGG GAC AAG TTT GTA CAA AAA AGC AGG CTT AGC CAC CAT GCC CCC TCT GCA CCA CG-3' |
| | | Reverse | 5'-GGG GAC CAC TTT GTA CAA GAA AGC TGG GTT CAC TCC TGC GTC CTC ATC GTC A-3' |
| *Homo sapiens* | Hs-EMRE-ct | Forward | 5'-GGG GAC AAG TTT GTA CAA AAA AGC AGG CTT AGC CAC CAT GGC GTC CGG AGC GGC-3' |
| | | Reverse | 5'-GGG GAC CAC TTT GTA CAA GAA AGC TGG GTT CCA GAA CTT CAG CCA GCC CC-3' |
| *Cyphellophora europaea* | Ce-MCU | Forward | 5'-GGG GAC AAG TTT GTA CAA AAA AGC AGG CTT AGC CAC CAT GAC TAA AGG CAA GCT GTT GAC GAC-3' |
| | | Reverse | 5'-GGG GAC CAC TTT GTA CAA GAA AGC TGG GTT TCT TGG TTC CGT TGT TGT TCT TTC GC-3' |
| *Fusarium graminearum* | Fg-MCU | Forward | 5'-GGG GAC AAG TTT GTA CAA AAA AGC AGG CTT AGC CAC CAT GAA CCA CGC TCT AAG GCG C-3' |
| | | Reverse | 5'-GGG GAC CAC TTT GTA CAA GAA AGC TGG GTT CGG CCA GGG CCG CAA-3' |
| *Metarhizium acridum* | Ma-MCU | Forward | 5'-GGG GAC AAG TTT GTA CAA AAA AGC AGG CTT AGC CAC CAT GGG CCA TGT CTT GGG TGG-3' |
| | | Reverse | 5'-GGG GAC CAC TTT GTA CAA GAA AGC TGG GTT GGT CCC AGC CCA TAT CGG TGT-3' |
| *Neosartorya fischeri* | Nf-MCU | Forward | 5'-GGG GAC AAG TTT GTA CAA AAA AGC AGG CTT AGC CAC CAT GCG GGC GCT TGT TAG CC-3' |
| | | Reverse | 5'-GGG GAC CAC TTT GTA CAA GAA AGC TGG GTT GCG CGT CAC ACT CAT GCT TGA-3' |

**Supplementary Table S3**. List of PCR primers used to amplify cDNAs from pLX304 vector to perform cloning into yeast expression vectors.

| Species | cDNA | Direction | Primer sequence |
|---|---|---|---|
| *Spizellomyces punctatus* | Sp-MCU | Forward | 5'- GGG GGA TCC ATG CGA ATC CCC TTC CA-3' |
| | Sp-fsMCU | Forward | 5'- GGG GGA TCC ATG GTG AGC CTG ATT CCA G-3' |
| | Sp-EMRE | Forward | 5'- GGG GGA TCC ATG TCC CGC ATT CTG ACA -3' |
| *Allomyces macrogynus* | Am-MCUa | Forward | 5'- GGG GGA TCC ATG CTG TCA AGG GCT CTG-3' |
| | Am-MCUb | Forward | 5'- GGG GGA TCC ATG CTG ATT TCT TGT CGC C-3' |
| | Am-fsMCU1 | Forward | 5'- GGG GGA TCC ATG TTC GCA CAG TCC CG-3' |
| | Am-EMRE1 | Forward | 5'- GGG GGA TCC ATG CCT CAG CTG CAT TTC T-3' |
| | Am-EMRE2 | Forward | 5'- GGG GGA TCC ATG CCC CCT CTG CAC-3' |
| *Cyphellophora europaea* | Ce-MCU | Forward | 5'-GGG GGA TCC ATG ACC AAG GGC AAG CT-3' |
| *Fusarium graminearum* | Fg-MCU | Forward | 5'-GGG GGA TCC ATG AAC CAC GCC CTG AG-3' |
| *Metarhizium acridum* | Ma-MCU | Forward | 5'-GGG GGA TCC ATG GGA CAC GTG CTG G-3' |
| *Neosartorya fischeri* | Nf-MCU | Forward | 5'-GGG GGA TCC ATG AGA GCC CTG GTG TCT-3' |
| *Homo sapiens* | Hs-EMRE | Forward | 5'-GGG CCC GGG ATG GCG TCC GGA GC-3' |
| | V5 | Reverse | 5'-GGG CTC GAG CTA CGT AGA ATC GAG ACC GAG-3' |
